# Restriction of intercellular communication is required for organ regeneration

**DOI:** 10.1101/2023.12.10.570601

**Authors:** Itay Cohen, Hagit Hak, Jessica Perez-Sancho, Ziv Spiegelman, Idan Efroni

## Abstract

The excision of the root tip, including the stem cell niche, triggers rapid regeneration from remnant cells in the stump. In plants, cell identity relies on positional information transported through cytoplasmatic bridges known as plasmodesmata. However, how such information is reset to allow the regeneration of lost identities is unknown. Here, we show that the movement of mobile signals is temporarily restricted near the incision site and that regeneration-induced members of the LATERAL ORGAN DOMAIN (LBD) plant-specific developmental regulators are necessary and sufficient for this restriction. Regeneration was disrupted in high-order *lbd* mutants but was restored by transient localized plasmodesmata closure. We propose that LBD-mediated modulation of intercellular connectivity is crucial for plant regeneration and may have widespread roles in *de novo* pattern formation.

**One Sentence Summary:** Plant-specific DNA binding genes mediate a transient restriction of intercellular communication to allow root regeneration

## Introduction

When the tip of the root, including its stem cell niche, is surgically removed, cells in a region near the cut site are rapidly recruited to regenerate the missing tissues (*1*). This process initiates with rapid activation of wound-induced genes and hormone biosynthesis (*2*–*4*), which causes differentiated cells to lose their old identity and enter a transitional mixed-identity state. During the following 48h, new patterns, including the *de novo* formation of the stem cell niche and missing distal tissues, are gradually established (*5*). As plant cells are immotile, this *de novo* pattern formation occurs within the context of an already patterned organ. While early stages in regeneration have been investigated in some detail, how *de novo* pattern formation occurs in later stages is unknown.

Cell identity in plants is primarily governed by a range of mobile signals, including proteins, RNAs, and hormones that travel through the symplastic route, made up of plasmodesmata connecting cytosols of adjacent cells (*6*–*9*). For example, the transcription factor SHORTROOT (SHR) is produced in the root internal stele and actively transported through plasmodesmata to the surrounding endodermis and quiescent center (QC) tissues, where it enters the nucleus (*10*–*12*). Reduced SHR activity or restriction of stele to endodermis movement results in loss of endodermal identity and disruption to cell division patterns (*13*–*17*). Other morphogenetic factors, such as the phytohormone auxin, whose spatial distribution instructs the position of the stem cell niche and distal root identities (*18*), also move through plasmodesmata (*19*). While auxin primarily moves through PIN-FORMED membrane-bound transporters (*20*), symplastic auxin transport plays a significant role during lateral root initiation (*21*), emergence (*22*), root tip regeneration (*3*), and vein patterning (*23*).

In some developmental scenarios, such as shoot meristem homeostasis (*24*–*26*), lateral root and nodule formation (*27*, *28*), vein patterning (*23*), and embryogenesis (*29*–*33*), coordinated reduction of plasmodesmata permeability forms semi-isolated regions (symplastic domains), with restricted connectivity to the surrounding tissue. Such regions can be formed by altering the number, direction, and structure of plasmodesmata (*34*, *35*) as well as by controlled deposition of β-1,3-D-glucan (callose) at the plasmodesmata neck (*32*). Callose restricts transport through plasmodesmata and is produced by plasmodesmata-localized callose synthetase genes (CalS), whose activity is modulated by PLASMODESMATA-LOCATED PROTEIN (PDLPs). Other, less characterized, pathways for regulating plasmodesmata permeability also exists (*6*). Symplastic domain are hypothesized to be involved in organ formation and growth regulation (*36*, *37*), but their developmental role and regulation is unclear

## Results

### Intercellular movement is transiently restricted during root regeneration

To map symplastic connectivity dynamics during root tip regeneration, we first characterized the movement of mobile signals during root tip regeneration. We used *SUC2:GFP* plants, in which GFP is produced in the proximal phloem and is allowed to freely diffuse in the meristem through the symplastic path (*38*) (Fig. 1A). In uncut roots, GFP signal is found throughout the root meristem, but in cut roots, GFP signal was gradually restricted from the distal part of the regenerating root and by 24h after the cut, was almost undetectable in a dome-like region, about ∼50µm from the cut site. By 72h post-cut, the signal was recovered at the distal part of the newly formed root tip (Fig. 1B-F). To verify that the loss of GFP signal at the tip is not due to effects on GFP fluorescence or SUC2 expression levels, we injected the inert symplastic dye HPTS at 1cm above the dissection site and followed the signal accumulation in the meristem at 3h after injection (Fig. 1G). In uncut roots, HPTS accumulated most rapidly in the cortex, consistent with the high apical-basal connectivity of this tissue (*34*). During regeneration, HPTS movement was restricted in a similar spatial and temporal pattern to GFP, confirming the inhibition of symplastic movement at the distal region (Fig. 1F-K). These observations are consistent with previous findings that, at 24h after the cut, stele-produced *SHR-GFP* fusion proteins could move laterally but did not move distally to the root end (*5*).

**Figure 1.**
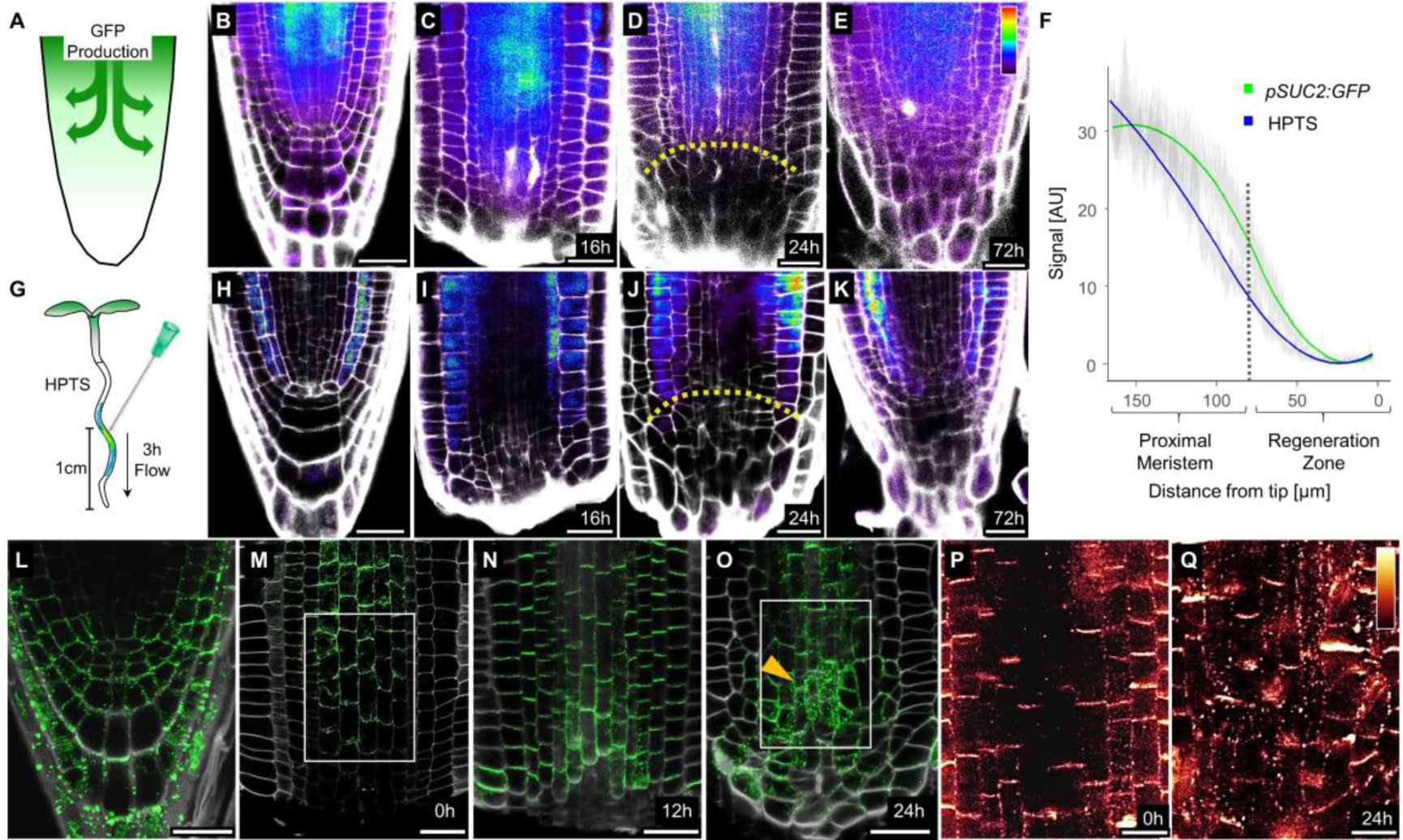
Transient and local symplastic movement restriction during regeneration. **A)** Illustration showing the *SUC2:GFP* movement assay. **B-E)** Confocal images of *SUC2:GFP* root meristems showing GFP intensity before (B) at 16h (C) 24h (D) and 72h (E) after the cut. **F)** Quantification of GFP or HPTS signal in regenerating meristems at 24h after the cut. Gray band is standard error (n=11). **G)** Illustration showing the HPTS injection assay. **H-K)** Confocal images of root meristems 3h after injection of HPTS before (H), at 16h (I), 24h (J) and 72h (K) after the cut. **L-O)** Confocal images of *35S:PDLP5-GFP* root tips before (L), at 0h (M), 12h (N), and 24h (O) after the cut. Arrowhead marks region of PDLP5-GFP accumulation. **P-Q)** Confocal images of anti-callose immunostaining of root at 0h (P) and 24h (Q) after the cut. Shown are the regions boxed in (M) and (O) respectively. Scale bars are 10µm (P-Q) and 25µm in (B-E, H-K and L-O).

The localized inhibition of symplastic movement suggests that plasmodesmata connectivity or permeability is altered during regeneration. To characterize plasmodesmata distribution, we followed the plasmodesmata marker PDLP5-GFP, expressed under a constitutive 35S promoter (*39*). PDLP5 induces callose deposition at plasmodesmata when strongly expressed (*39*), but weak expression lines can be used to visualize plasmodesmata with minimal effects (*22*). Before the cut, a punctate pattern of PDLP5-GFP, consistent with its plasmodesmata localization, had a strong signal at the columella and lateral root cap (Fig. 1L). Immediately after and until 16h after the cut, PDLP5-GFP was localized to plasmodesmata-rich proximal-distal cell walls (*34*) (Fig. 1M-N), but at 24h post-cut, there was a marked change in PDLP5-GFP signal, and it accumulated in a central distal region, up to ∼80µM from the cut tip (Fig. 1O). Anti-callose immunostaining showed that, together with the PDLP5-GFP signal, callose significantly accumulated in a punctate pattern at internal root tissues at 24h after the cut, as compared to immediately after injury (Fig. 1P-Q; Fig. S1A-C). These observations were consistent with the inhibition of GFP and HPTS movement and indicate that, during regeneration, plasmodesmata permeability is dynamically altered to isolate a distal region of the root. Notably, this region is where the new stem cells and distal identities are specified at 16h to 48h after the cut (*5*).

### Regeneration-induced GATEKEEPER genes are required for symplastic movement restriction

The presence of temporarily isolated symplastic domains was shown to rely on callose metabolic enzymes (*40*), but its’ developmental regulators remain unknown. To identify the regulators of movement inhibition during regeneration, we used single-cell transcriptional profiles to identify factors upregulated at the distal part of the regenerating roots (*5*). We noticed that five root-expressed LATERAL ORGAN BOUNDARY DOMAIN/ASYMETRIC LEAVES 2-LIKE genes (*LBD6/AS2*, *LBD12*, *LBD13*, *LBD15* and *LBD36*) were co-induced during regeneration in a transient pattern (Fig. S2A-B). Promoter fusion for *LBD36* confirmed the single-cell derived expression pattern and the induction of this gene near the cut site (Fig. S2C-G). LBD/ASL is a family of plant-specific DNA-binding, nucleus-localized proteins associated with organ boundary formation (*41*), and regulating many developmental processes, including root initiation and emergence, cambium growth, leaf dorsiventrality, vein patterning, embryo development, inflorescence branching and callus formation (*42*–*51*).

To determine whether the induced LBD genes are responsible for the symplastic isolation of the regenerating root tip, we used CRISPR to generate high-order mutants and tested at least two alleles for each gene (Table S1). Single *lbd6* and *lbd36* had no observed developmental phenotype in the root, as previously reported (*52*). However, high-order mutants displayed a gradual loss of organization at the distal part of the root meristem and progressively shorter roots (Fig. 2A-E; Fig. S3A). The quadruple *lbd6 lbd36 lbd12 lbd15* also had a reduction in the number of root hairs and a delay in lateral root emergence (Fig. S3A-C). These phenotypes were consistent with the genes root cap and epidermis expression patterns (Fig. S2B). We could not obtain viable *lbd13* mutants, as was previously reported (*53*), and thus focused on the quadruple mutant (referred to as *quad*).

**Figure 2.**
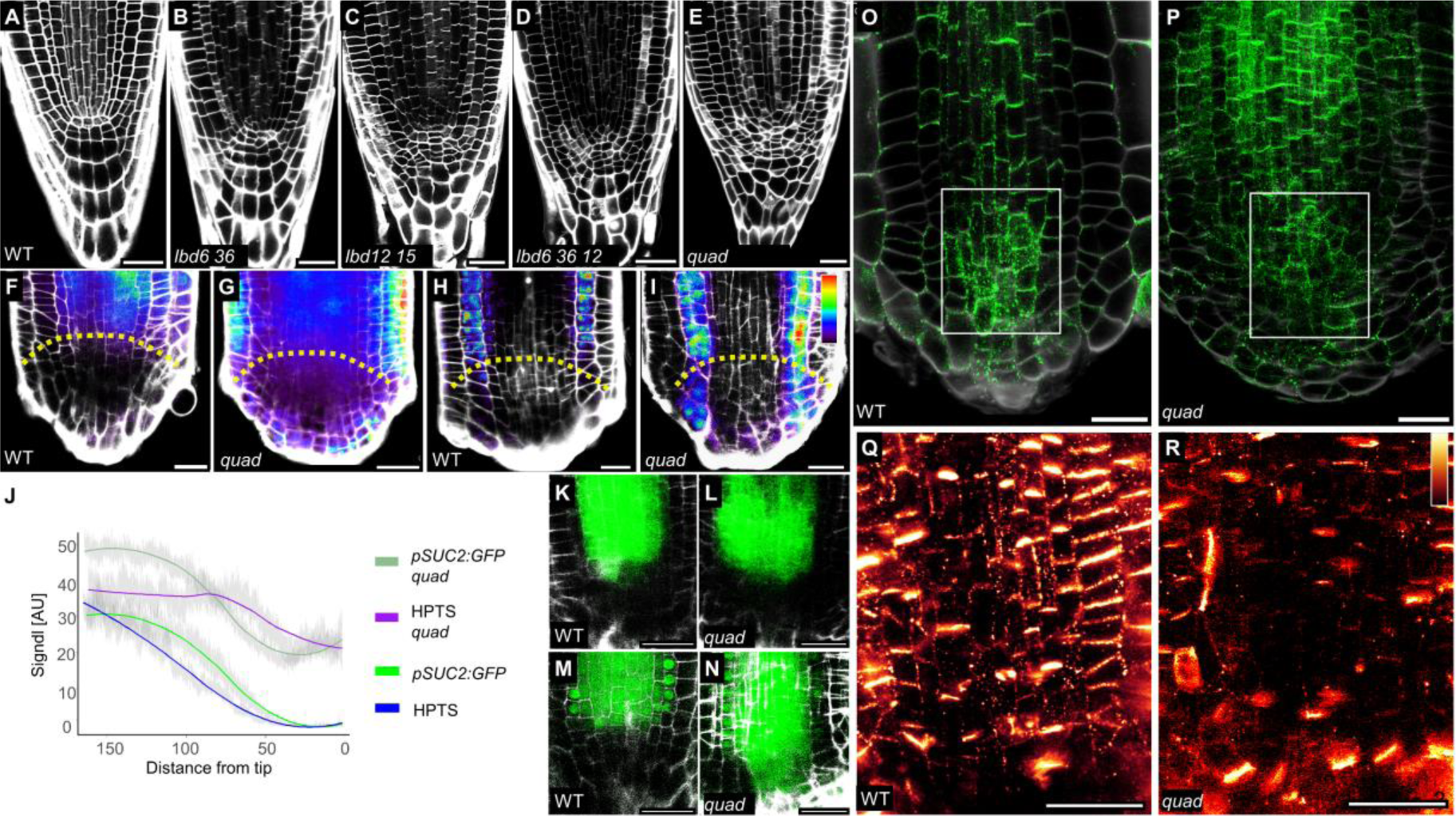
GATEKEEPER genes are required for symplastic isolation during regeneration. **A-E)** Confocal images of primary root tips of 7-days old seedlings. **F-I)** Confocal images of *SUC2:GFP* (F-G) and HPTS-injected (H-I) in WT (F,H) or *quad* (G,I) background. Dotted line marks the region of symplastic movement restriction in WT. **J)** Quantification of the signal in (F-I). Gray band is standard error (n>10). **K-N)** Confocal images of root tips at 24h after cut of *pSHR:ER-GFP* (K-L) and *pSHR:SHR-GFP* (M-N) in WT (K,M) and *quad* (L,N) backgrounds. **O-R)** Confocal images of root meristems at 24h after the cut of *35S:PDLP5-GFP* plants (O-P) or immunostained using anti-callose antibodies (Q-R) in WT (O, Q) or *quad* (P, R) background. Rectangle in O, P mark the regions shown in Q, R, respectively. Scale bars are 25µm.

When the *SUC2:GFP* reporter was introduced into the *quad* background, free mobile GFP accumulated at the distal zone of regenerating roots, suggesting little restriction on symplastic movement, an observation we validated using injected HPTS (Fig. 2F-J). Furthermore, while the expression of the transcriptional reporter *pSHR:ER-GFP* was excluded from the distal zone in both WT and the *quad* at 24h after the cut, the *pSHR:SHR-GFP* protein signal was detected in the distal zone of mutant plants (Fig. 2K-N). At 12h after the cut, there was no marked difference in PDLP5-GFP signal between the wild type and *quad* roots (Fig. S3D-G). However, by 24h after the cut, and consistent with the loss of movement restriction, *quad* roots failed to accumulate PDLP5-GFP signal at the distal part of the root, and there was a significant reduction in callose deposition (Fig. 2O-R; Fig. S4A-C), indicating that *quad* mutant plants fail to establish symplastic isolation during regeneration. This effect was local to the distal part of the root, as PDLP5-GFP signal increased at proximal regions of *quad* mutant roots.

We conclude that the four tested LBDs are required for restricting movement near the cut site during regeneration. These co-regulated LBDs are not phylogenetically related and are classified as classes IA2, IC1, and IC2 (*54*). Due to their common effect on symplastic transport, we collectively refer to them as GATEKEEPER genes.

### Ectopic expression of GATEKEEPER genes restricts symplastic movement

To test whether the GATEKEEPER genes are sufficient to restrict symplastic transport, we used a β-estradiol induction system to ectopically express *LBD6/AS2* and *LBD36/ASL1* in *SUC2:GFP* plants. Induced expression of either gene restricted GFP movement, a result we verified using injected HPTS (Fig. 3A-G). Similar restriction of injected HPTS movement was observed when *LBD12* or *LBD15* were overexpressed (Fig. S5A-C). Prolonged induction of all LBDs resulted in similar arrest of root growth and mitotic cell division, along with increased cell expansion (Fig. S5D-I), suggesting that in this context, all four LBDs can perform a similar function. To determine whether LBD overexpression can also inhibit directed movement, we tested the effect of *LBD36* transient expression on *pSHR:SHR:GFP* plants. After 6h of *LBD36* induction with β-estradiol treatment, SHR-GFP signal was lost in sporadic endodermal cells, and by 18h, ectopic lateral divisions of the endodermis were observed, a phenotype typical for low SHR levels (*17*) (Fig. 3H-K).

**Figure 3.**
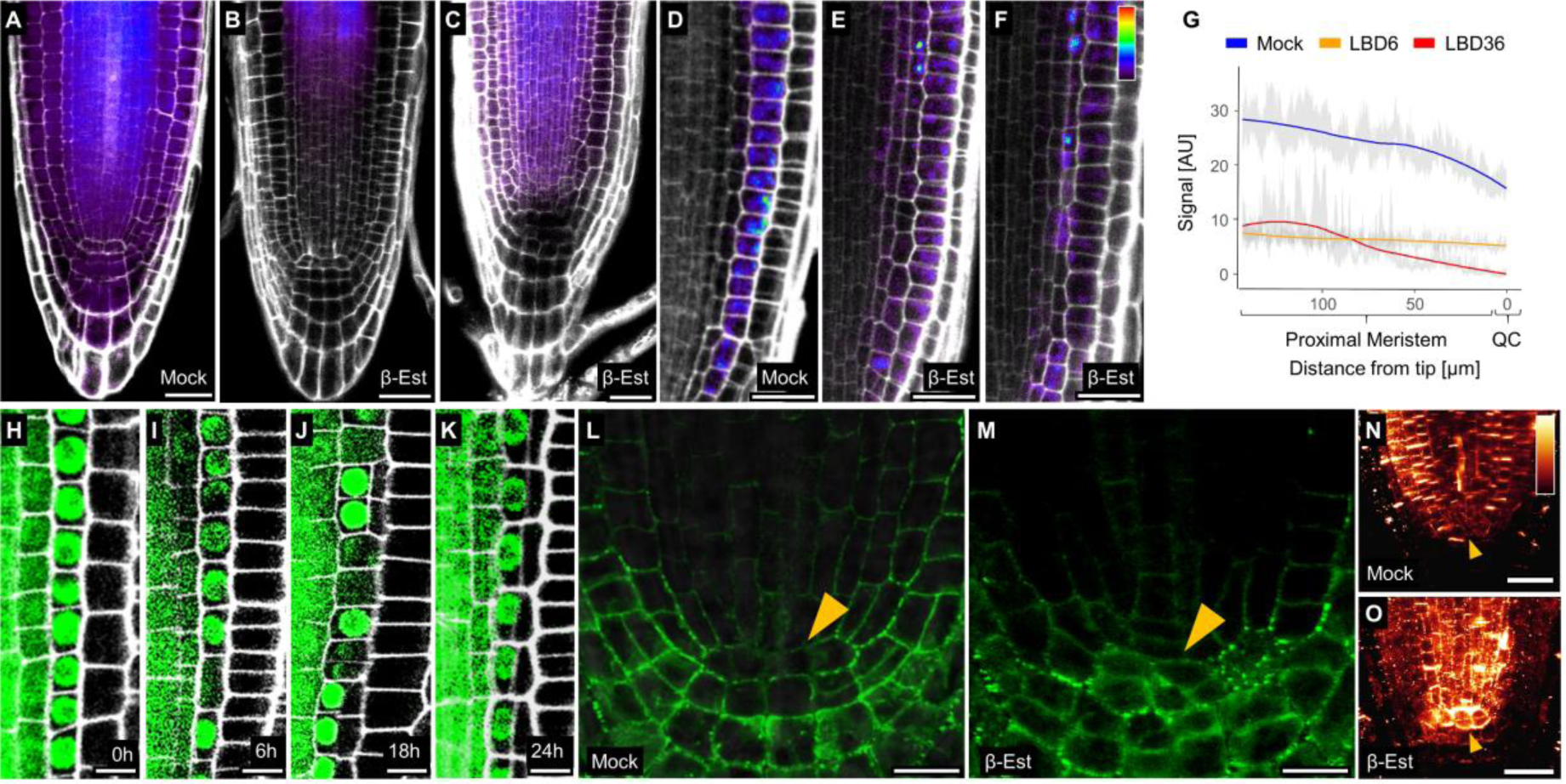
Ectopic expression of GATEKEEPER genes restricts intercellular movement. **A-F)** Confocal images of *SUC2:GFP* (A-C) and injected HPTS (D-F) *35:XVE lexA:LBD6* (B,E) and *35S:XVE lexA:LBD36* (A,C-D,F). Either mock treated (A,D) or treated with β-estradiol for 48h (B-C, E-F). **G)** Quantification of the HPTS signal in (D-F). Gray band is standard error (n>10). **H-K)** Confocal images of *pSHR:SHR-GFP 35S:XVE lexA:LBD36* plants immediately after (H), at 6h (I), 18h (J) and 24h (K) after treatment with β-estradiol. **L-M)** Confocal images of root meristem of *35:XVE lexA:LBD36 35S:PDLP5-GFP* plants treated with mock (L) or β-estradiol (M) for 24h. **N-O)** Immunostaining using anti-callose antibodies of *35:XVE lexA:LBD36* treated with mock (N) and β-estradiol for 24h (O). Arrowhead points to the position of the QC. Scale bars are 25µm (A-F, N-O) and 10µm (H-M).

Consistent with their restrictive effect on symplastic movement, induced overexpression of *LBD36* resulted in increased accumulation of PDLP5-GFP signal and increased callose staining, mainly in the stem cell region of the uncut root (Fig. 3L-O). To test the generality of the GATEKEEPER LBD, we transiently expressed *LBD6* and *LBD36* in the heterologous *Nicotiana benthamiana* leaf epidermal cell system. Aniline blue staining showed a marked increase in callose accumulation, suggesting a broad capacity for these LBDs to block plasmodesmata (Fig. S6A-E).

GATEKEEPER genes are expressed throughout development, and the loss of distal meristem cellular organization suggests that they may play a function in the regulation of symplastic transport outside the context of regeneration (Fig. 2A-E). Indeed, in uncut *quad* roots, there was reduced accumulation of PDLP5-GFP and callose staining, in particular at the distal part of the stem cell niche (Fig. S3H-K). Injection of HPTS into uncut roots revealed higher levels of accumulation at the distal part of the *quad* root (Fig. S3L-M). Further, SHR-GFP signal could be ectopically detected at the root cap of *quad* mutants, although we cannot rule out that part of this signal expansion is due to transcriptional effects, as expression of the *SHR* promoter was also altered in uncut mutant roots (Fig. S3N-Q).

Taken together, we show that *GATEKEEPER* genes are required and sufficient to enact symplastic transport restriction at the distal part of the root during later stages of regeneration and other developmental contexts.

### Tissue patterning is disrupted in regenerating roots of *gatekeeper* mutants

We next tested whether root regeneration is affected in *gatekeeper* mutants. Double mutants had normal regeneration rates. However, *lbd6 lbd36 lbd12* mutants and *quads* had a significant reduction in regeneration rates (ANOVA; p<0.05), although meristem size was not altered in any of the mutants (Fig. 4A-B). When root regeneration is disrupted by inhibition of wound-responsive genes or auxin biosynthesis, the meristem differentiates within 24h (*2*, *3*). However, analysis of regeneration in *quad* mutants showed that early stages of regeneration were similar to WT. By 24h after the cut, abnormal cell division patterns were observed at the distal part of the stump, and only by 48h-72h, the meristem began to differentiate (Fig. 4C-L), suggesting that *quad* mutants are defective in late patterning stages, rather than in early wound response.

**Figure 4.**
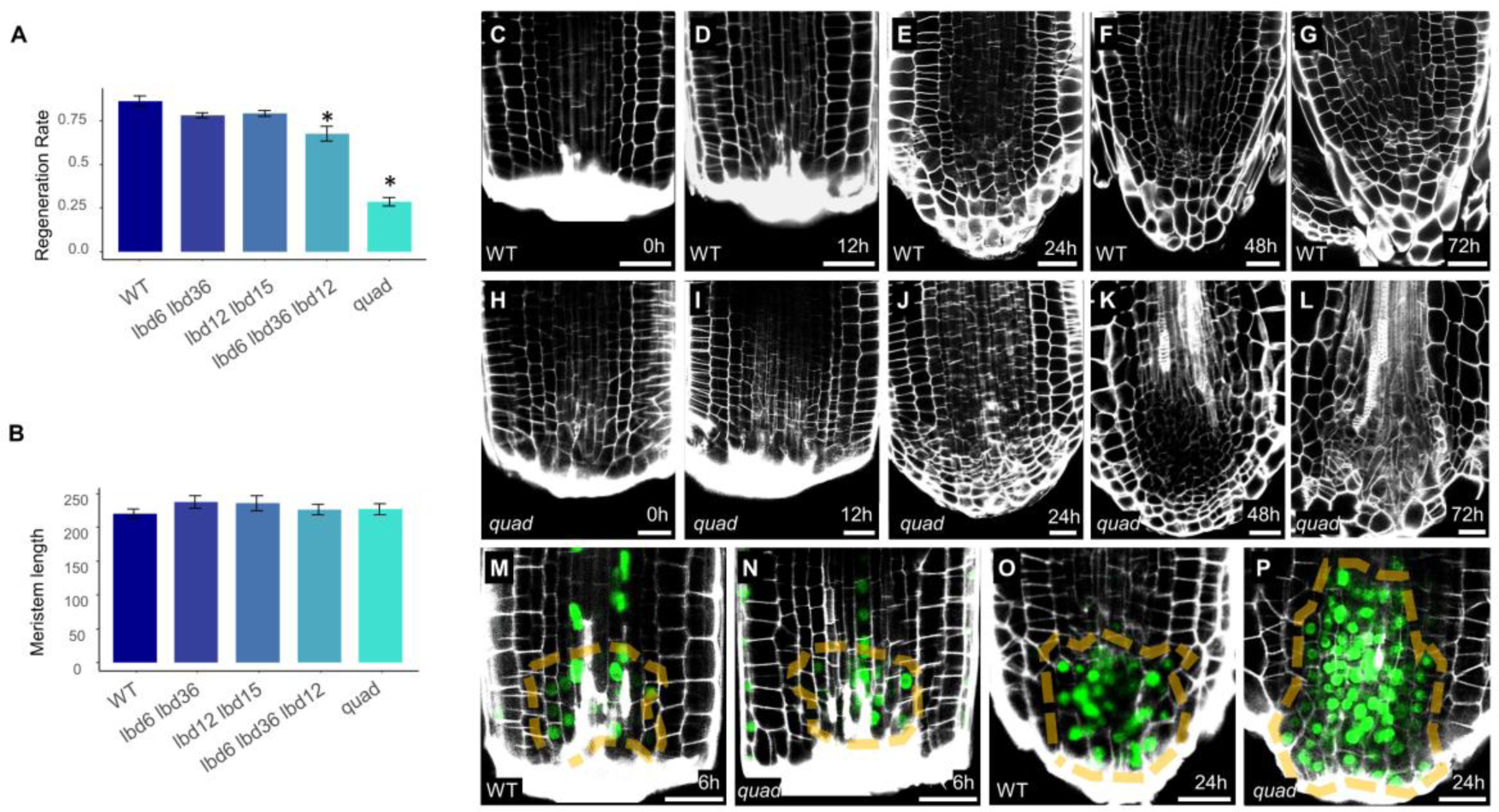
GATEKEEPER genes are required for patterning stage of root tip regeneration. **A-B)** Regeneration rates (A) and meristem lengths (B) for GATEKEEPER genes mutant combinations. Bars are standard error (n=3). Asterix indicate statistically significant difference from wild type (ANOVA; p<0.05) **C-L)** Confocal images of regenerating WT (C-G) and *quad* (H-L) root tips. **M-P)** Confocal images of *DR5rev:3xVENUS-N7* in the background of WT (M,O) or *quad* (N,P) at 6h (M-N) and 24h (O-P) after the cut. Dashed lines mark the auxin peak. Scale bars are 25µm.

To verify this observation, we followed the expression of the auxin response reporter and early-regeneration marker DR5, which is rapidly induced near the cut site due to local auxin biosynthesis (*3*). DR5 expression pattern in regenerating roots at 6h after the cut was similar between WT and *quad* (Fig. 4M-N). However, by 24h, the DR5 signal was diffuse and expressed throughout the stele, unlike the focused distal peak in the WT (Fig. 4O-P). Auxin is required for regeneration and *LBD6* was previously suggested to act upstream of the PIN auxin transporters (*3*, *52*). We therefore tested whether the regeneration defect results from low auxin levels by treating regenerating roots with either auxin or the auxin transport inhibitor 1-Naphthylacetic acid (NPA). However, neither auxin nor NPA could rescue regeneration in the mutant (Fig. S7A), and the effect of NPA on auxin distribution was additive (Fig. S7B-C), indicating that the failure to regenerate is not solely due to low auxin levels or transport defects.

To better characterize the regeneration defect in *quad* roots, we generated single-cell RNA-Seq profiles of WT and mutant meristems at 16h after the cut, after early wound responses, but before the appearance of morphological differences. Two replicates were collected from each genotype and combined with a reference single-cell dataset of an intact root meristem (*55*). Clusters were annotated based on Index of Cell Identity scores (*56*) and published tissue markers from high-resolution analyses (*57*) (Fig. 5A; Fig. S8 and S9).

**Figure 5.**
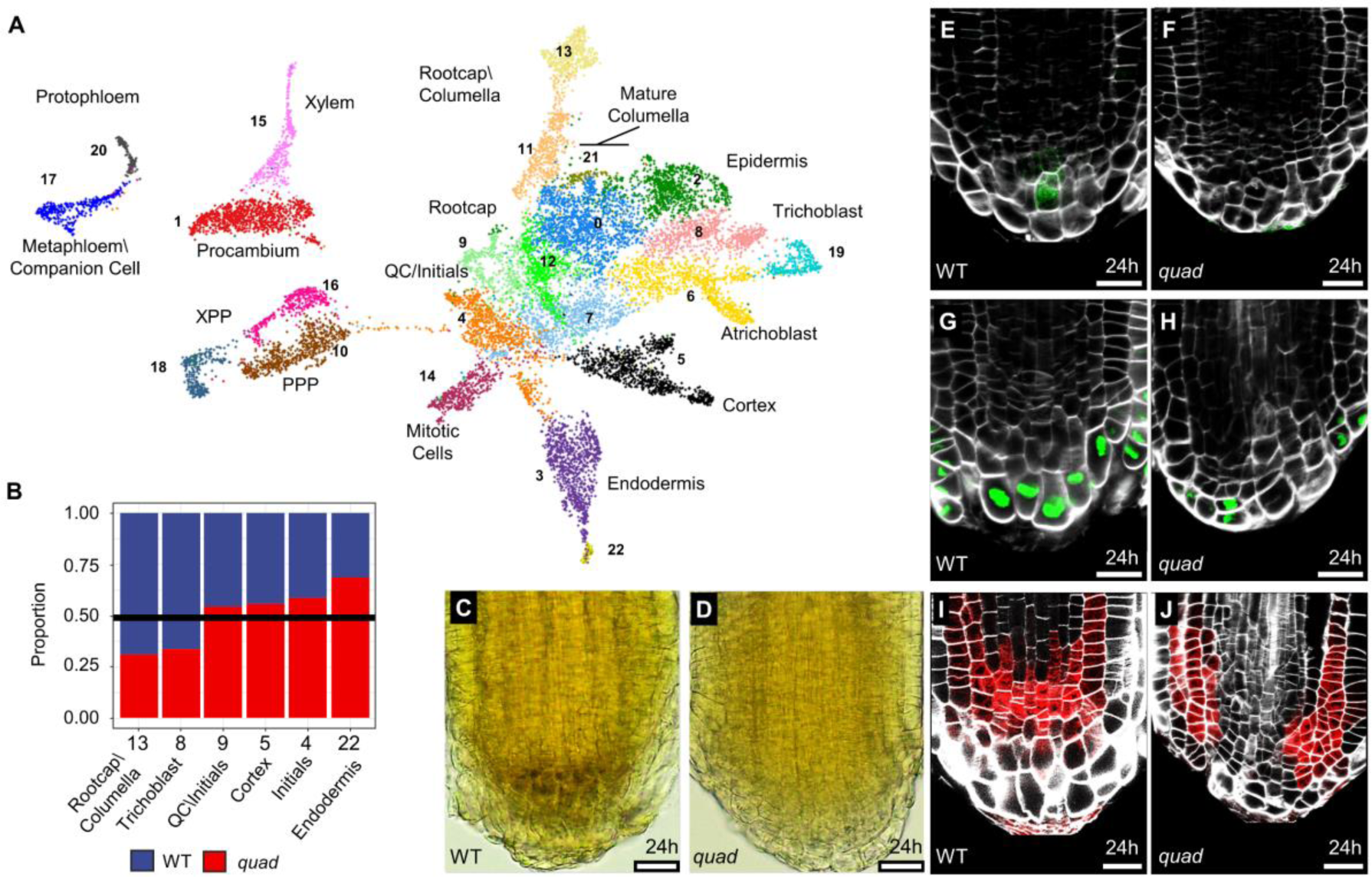
*gatekeeper* mutants are defective in specification of distal identities patterns. **A)** Combined UMAP of single-cell profiles from 16h after cut of WT and *quad,* and uncut WT root meristem. **B)** Relative representation of WT and *quad* cells in each cluster. **C-D)** Iodine staining showing starch accumulation in WT (C) and *quad* (D) at 24h after cut. **E-J)** Confocal images of regenerating roots of the late columella marker *PET111:GFP* (E-F), *SMB:GFP-NLS* (G-H) and *WOX5:mCherry* (I-J) in WT (E,G,I) and *quad* (F,E,J) backgrounds. Scale bars are 25µm.

All root identities were represented in the reference, wild type, and *quad* samples, with the exception of cluster 21, a subpopulation of mature root cap cells that were not found in the regenerating root samples (Fig. S10A-B). As expected, distal identities, which are excised in regenerating roots were underrepresented in regenerating roots, while internal tissues, such as procambium and vasculature tissue, were overrepresented (Fig. S10B). Cells in two clusters were underrepresented in *quad* mutants compared to wild type: trichoblasts (root hair cells) and rootcap/columella (χ-test, p<1E-4; Fig. 5B). Reduction in trichoblasts number was consistent with the observed reduced root hairs in the mutant (Fig. S3B-C), but a reduction in the number of newly specified rootcap/columella cells was unexpected. To verify this result, we examined the accumulation of starch-containing amyloplasts, which are characteristic of mature distal columella cells. As previously reported, amyloplasts were detected in wild type roots at 24h post-cut (*1*). However, they were almost entirely missing from the distal end of the regenerating *quad* roots. In agreement, expression of the mature columella marker PET111 was absent in the mutant at the same timepoint, indicating a defect in distal identity acquisition (Fig. 5C-F).

To determine the spatial defect in distal identity patterning in *quad* mutants, we analyzed the expression of the early root cap reporter *pSMB:mNeonGreen* (*58*). In regenerating wild type roots at 24h after the cut, *pSMB:mNeonGreen* marks the entire distal layer (Fig. 5G). SMB-expressing cells were specified in the mutant, but their expression was excluded from the center of the root (Fig. 5H), suggesting the central region of the root cannot correctly specify the new tissues. To determine the extent of the patterning defect, we also examined the QC reporter *WOX5:mCherry* which is induced early in regeneration and, at about ∼24h, is found in a broad central region next to the cut site (*1*, *5*) (Fig. 5I). *WOX5:mCherry* was expressed in *quad* mutants, but, consistent with the defect in *SMB* patterning, it was excluded from the central region of the root (Fig. 5J). These results indicate that distal identities typical of regeneration are specified in the mutant, but their spatial pattern is disrupted at a stage when the distal region of the root is normally symplastically isolated.

### Restricting symplastic transport restores regeneration to *gatekeeper* mutants

We then asked whether we could restore regeneration capacity to *quad* mutants if we enact transient symplastic isolation at the newly specified distal cells. We generated plants where *PDLP5*, which restricts symplastic transport when strongly overexpressed (*39*), is expressed under an inducible promoter of the GATEKEEPER gene *LBD36*. To transiently induce symplastic isolation, we cut root meristems, allowed them to recover for 6h, induced PDLP5 expression for 18h using β-estradiol, and then moved them back to recovery media (Fig. 6A). As expected, induced expression of *PDLP5* inhibited the movement of the symplastic tracer HPTS into the *quad* mutant distal zone (Fig. 6B-C). Transient induction of *PDLP5* did not affect regeneration of wild type plants but, remarkably, could significantly recover regeneration, distal patterning, and starch accumulation in *quad* mutants (P-val <0.01; n=3; Fig. 6D-H). We conclude that restriction of symplastic transport is required for proper pattern acquisition and is sufficient to rescue both regeneration and patterning in *quad* mutants.

**Figure 6.**
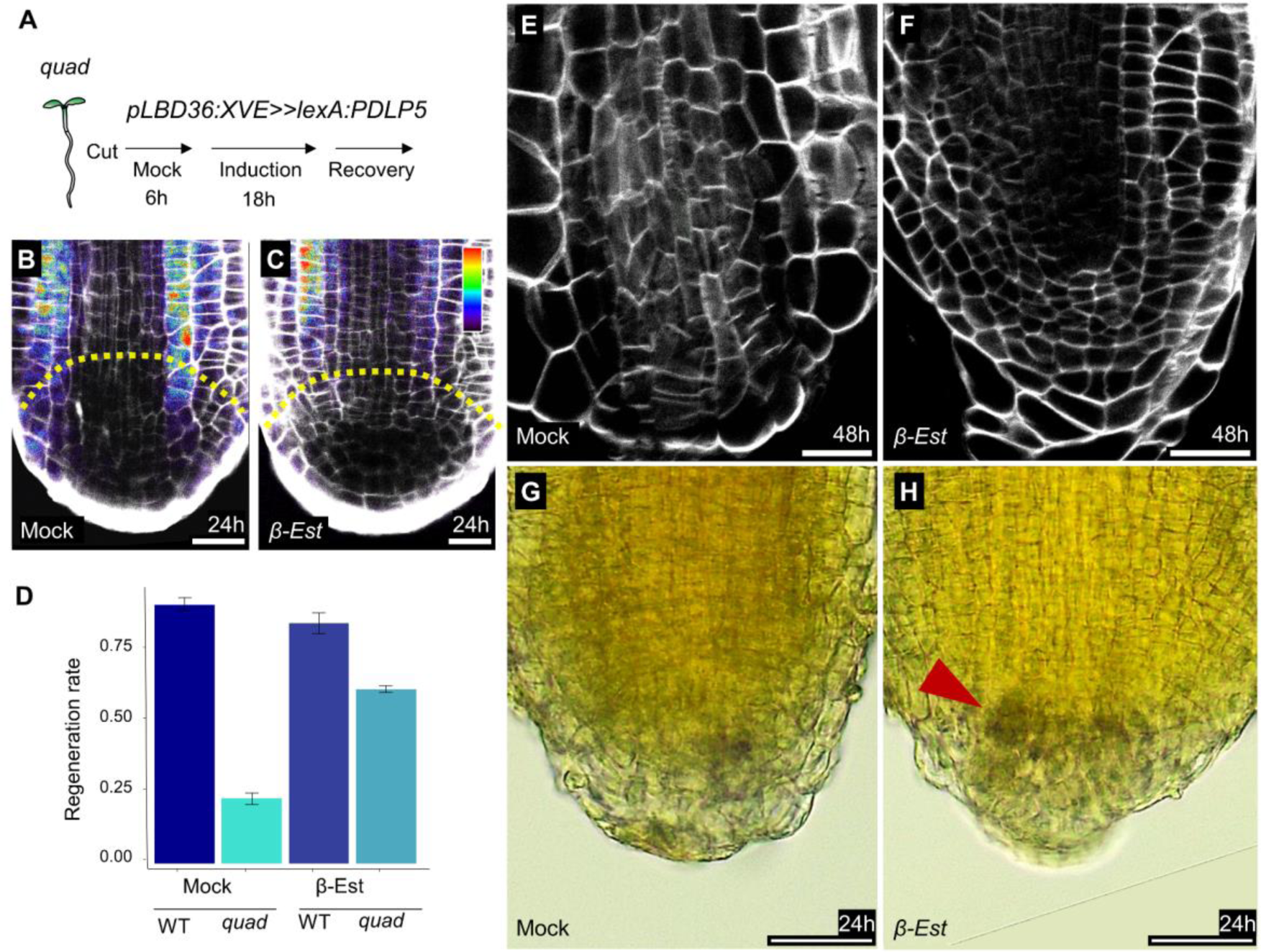
Transient restriction of intercellular connectivity in *gatekeeper* mutants. **A)** Illustration showing transient restriction protocol. **B-C)** Confocal images of *pLBD36:XVE lexA:PDLP5* root meristems at 24h after injection with HPTS following treatment with mock (B) or induced with *β-estadiol* (C) for 18h. **D)** Regeneration rates of WT and *pLBD36:XVE lexA:PDLP5* plants. **E-H)** Confocal images (E-F) or iodine starch staining (G-H) of *pLBD36:XVE lexA:PDLP5* plants at 24h (G-H) or 48h (E-F) after the cut. Red arrowheads point to increased starch accumulation in b-estradiol treated plants. Scale bars are 25µm.

## Discussion

Plants possess a remarkable ability to regenerate from severe damage. We show that this process requires the formation of a symplastic domain at the patterning stage of regeneration. Importantly, this domain is not required for the acquisition of specific identities, indicating that the morphogenetic factors that specify cell fates are present from an early stage of regeneration. However, the symplastic domain is essential for establishing and stabilizing multicellular patterns within the regenerating organ, possibly by limiting the diffusion of the morphogenetic factors, such as auxin. The presence of such diffusion limiting domain also occurs during the formation of somatic embryos in tissue culture (*33*), and may be common to many instances of *de novo* pattern formation.

Our work shows that the symplastic domain is produced by a subset of LBD genes. Domain swapping and overexpression experiments showed that different LBDs have, at least in part, distinct functions (*52*, *59*, *60*). The GATEKEEPER LBDs are not phylogenetically related, suggesting that while some members may have distinct functions, the regulation of symplastic transport could be shared within the LBD family. Consistent with this hypothesis, the activity of LBD genes is required for multiple developmental processes involving the transient formation of symplastic domains, such as lateral root formation and emergence (*22*, *27*, *61*), nodule formation (*28*, *61*, *62*), and vein patterning (*23*). LOB, the founding member of the LBD group is expressed organ boundaries, as are other members of this family (*42*, *50*). Thus, LBD-mediated restriction of morphogenic factor mobility may be a crucial part of organ formation in response to wounding as well as during iterative plant development.

## Acknowledgments

We thank E. Bayer, Y. Eshed, D. Jackson, and S. Savaldi Goldstein for helpful comments and discussions. J.Y. Lee for providing materials.

## Author contributions

Conceptualization: IC, IE and ZS

Methodology: IC, JPS and IE

Investigation: IC and HH

Supervision: IE

Writing: IC and IE

## Data and material availability

Raw and processed single-cell data is available in GEO series GSE248204. Biological material available from the authors by request.

## Materials and Methods

### Plant materials and growth conditions

Arabidopsis seeds were surface sterilized by chlorine gas treatment (5ml HCl added to 95ml of bleach enclosed in an airtight container) for 2h. After sterilization, seeds were imbibed in sterile DDW before sown on ½ Murashige and Skoog (Sigma) medium (2.2gr/l MS, 0.5% sucrose, 0.8% plant agar (Duchefa), pH 5.7) containing plates. After 2 days of stratification at 4°C, plates were placed vertically at 16h light (120 μE m^−2^sec^−1^) and 8h dark cycles at 22°C. Unless noted otherwise, 6-day-old seedlings were used for regeneration, microscopy, and RNAseq assays. *pSUC2:GFP, pSHR:SHR-GFP, pSHR:ER-GFP, pDR5:3xVENUS-N7, pWOX5:ER-mCherry, WIP4:ER-GFP* were previously published(*5*, *38*, *63*–*65*).

### Transgenic lines

Transgenic lines were generated using the MoClo Golden Gate system (*66*) with domesticated assembled fragments. Synthesized DNA fragments (via Syntezza Bioscience Ltd.) were altered to remove BpiI\BsaI restriction sites. The promoter of LBD36 (1,849bp upstream to the translation start site; pLBD36) was cloned into pICH41295. The terminator of LBD36 (1,127bp downstream to the stop codon; LBD36^ter^) was cloned into pICH41276. The pLBD36:NeonGreen-N7:LBD36ter reporter line was assembled from LBD36^ter^, mNeonGreen (synthesized and cloned into pAGM1299), N7 nuclear localization signal in pAGM1301 (as previously described (*3*) and terminator, into the level 1 plasmid pICH47742. Inducible overexpression (OE) constructs were based on the XVE system (*67*). The constitutive promoter 35SX2 was cloned into pICH41295. LexA-VP16 and hER were cloned into pAGM1287 and pAGM1301, respectively. The rbcS-E9 terminator was cloned into pICH41276. The 35SX2:LexA-VP16-hER:rbcS-E9t was assembled into the level 1 plasmid pICH47742. LBD GATEKEEPER genes CDS regions (LBD6 and 36 were amplified from Col-0 cDNA; LBD12 and 15 were synthesized) were cloned into pICH41308 and assembled downstream to pLexA (in pICH41295) and upstream to the NOS terminator (in pICH41276), into the level 1 plasmid pICH47751 to generate the plexA:GATEKEEPER_LBD_CDS:NOSter construct. For PDLP5-SlNeonGreen inducible expression construct, pLBD36 in pICH41295 assembled (upstream to the previously described LexA-VP16-hER and the NOS terminator) into the level 1 backbone pICH47742 to construct pLBD36: LexA-VP16-hER:rbcS-E9ter. PDLP5 CDS region (minus the stop codon) was synthesized and cloned into pAGM1287. mNeonGreen (synthesized and cloned into pAGM1301) was oriented to enable a transcriptional fusion with PDLP5. plexA:PDLP5-mNeonGreen:OCSter was assembled from the lexA operator, the fused PDLP5-mNeonGreen, and the OCS terminator (synthesized and cloned into pICH41276), which were collectively inserted into the level 1 pICH47751 backbone. The *35S:PDLP5-GFP* construct was kindly provided by Jung-Youn Lee. Binary vectors (pAGM4723 as the backbone) T-DNA regions carried the constitutively (35S) expression unit of neomycin phosphotransferase II (NPTII) as a selectable marker, except for *PDLP5-SlNeonGreen*. Binary vectors were introduced into Columbia-0 ecotype plants, or the *quad* mutants by floral dipping using agrobacterium tumefaciens strain GV3101. Primers are provided in Supplemental Table S3.

### CRISPR mutants

To generate CRISPR/Cas9 alleles, sgRNA were cloned into level 1 backbones downstream to the AtU6 promoter, as previously described(*46*), with each gene targeted with two guides. sgRNA sequences were designed using the Benchling CRISPR gRNA design tool (http://www.benchling.com) and were cloned into a pAtUBQ10:Cas9 expressing backbone (based on pAGM4723) 57. Two constructs were built: 1) to target LBD6 and LBD36, 4 guides were cloned into level 1 plasmids pICH47761, pICH47772, pICH47781 and pICH47791. 2) To target all five GATEKEEPER LBDs, the first construct (level 2) was re-opened, allowing for an additional 6 guides (in plasmids pICH47732, pICH47742, pICH47751, pICH47761, pICH47772, and pICH47781) to be cloned into it. For genotyping, genomic DNA was extracted by grinding ∼10mg of leaf tissues in sucrose solution (50mM Tris-HCl; pH7.5, 300 mM NaCl, 300mM sucrose) followed by 100C° for 15 min. Diluted (1:4) supernatant of the extracts was used for PCR (2xPCRBIOTaqMixRed, Biosystems Company) and amplified using designated primers (Table. S2). Genotypes were verified using electrophoresis and Sanger sequencing of the amplified fragments.

### Regeneration, movement assay, and starch staining

For the root rip regeneration assay, 6-day-old seedlings were excised at ∼130 µm from the root tip using a dental needle, as described previously(*1*). At least three biological replicates with at least 50 plants/replicate were used. For tracing of symplastic movement, 8-Hydroxypyrene-1,3,6-trisulfonic acid trisodium salt (HPTS; Sigma; 5mg/ml in 0.4% plant agar) was introduced by performing a minor wound to the epidermis with a fine dental needle 1cm from the root tip. The dye solution was then pipetted directly on the wound. HPTS migration towards the meristem was enabled, while plates were kept in the dark for ∼4h prior to their imaging. For starch-containing amyloplasts of mature columella cells, seedlings were incubated in Lugol’s Iodine solution (Sigma-Aldrich) for 30 seconds and then washed with DDW and mounted with chloral hydrate (water: chloral hydrate: glycerol, 8∶2∶0.25, v/w/v).

### Microscopy and phenotype characterization

Confocal imaging was performed using a Leica SP8 confocal microscope. Cellular borders were visualized using Propidium Iodide solution (0.01mg/ml, Sigma) for live samples or SCRI Renaissance 2200 (SR2200, 0.1% (v/v)) for fixed and cleared samples, using the ClearSee protocol(*68*). Fluorescent signal intensity measurements were obtained via ImageJ. The mean gray value profile of each image was collected from the cortex file for the HPTS signal or the middle of the root for the *pSUC2:GFP* signal. Nikon SMZ18 fluorescent stereomicroscope was used for standard visualization and imaging of fluorescent markers expression and root phenotypes. A Leica ICC50W light microscope was used for Lugol staining imaging.

### Callose immunolocalization and quantification

Immunolocalization of callose was carried out as previously described (*69*). Briefly, seedlings were fixed in 4% paraformaldehyde (Sigma), adhered to Polysine-coated slides (Thermo Scientific), followed by cell-wall digestion in 2% Driselase (GlpBio) at pH=5(*70*). (1-3)-β-glucan-directed monoclonal (Biosupplies Australia) diluted to 1:125 was used as primary antibody and Donkey Anti-Mouse IgG H&L Alexa Fluor® 555(Abcam) diluted to 1:500 was used as secondary. To quantify callose signal, roots were imaged at identical confocal settings. Contrast was adjusted to remove background signal and the average signal intensity at the central root region, between the endodermis layers, was measured using ImageJ. Signal from cell plates was manually removed from the measured region.

### Callose induction assay in *N. benthamiana* leaves

Agroinfiltration was performed as previously described (https://doi.org/10.1093/pcp/pcab179). Briefly, *Agrobacterium tumefaciens* strain EHA105 containing the *p35S:LBD6* and *p35S:LBD36* and the negative control *p35S:RBSC* (Rubisco) binary vectors were grown overnight at 28°C. Cultures were resuspended in 2-(N-morpholino)-ethanesulfonic acid (MES) buffer (10 mM MgCl_2_, 10 mM MES, and 150 µM acetosyringone, pH 5.6) to an optical density of OD_600_ = 0.25 and infiltrated using a needleless syringe into the abaxial side of the fourth or fifth leaf of 5-week-old *N. benthamiana* plants. Plants were monitored 24h post infiltration. Aniline blue staining was done as described by Hak et al. 2023 (https://doi.org/10.1111/mpp.13318). Aniline blue solution (0.1% in water) was infiltrated into the abaxial side of the leaf. Ten minutes later, leaf epidermal cells were imaged using an Olympus IX 81 inverted laser scanning confocal microscope (Fluoview 500). Excitation of aniline blue at 405 nm was done using a BA430–460 filter with a UPlanSApo 60× 1.35 oil objective, 8/0.17 FN26.5. Quantification was performed by measuring the aniline blue fluorescent signal in at least 15 fields from 3 individual leaves using the ImageJ software (https://imagej.nih.gov/ij/).

### Single Cell mRNAseq

For 10X genomics sequencing, ∼100 regenerating roots at 16hpc were harvested for each sample. Primary roots tips (∼1cm) were collected into a 35X10mm petri dish containing 60µl of cell wall digestion solution (3% w/v Cellulase, 1%w/v Macerozyme (Yakult), 0.4M Mannitol, 20mM MES, 0.02M KCl, Tris-HCl pH=5.7, heated to 55C° for 10min, cooled to room temperature followed by addition of 0.001 gr/ml BSA and 20mM CaCl2). Meristems were collected after 15 minutes of incubation and transferred to a fresh digestion solution for 1h. The solution was filtered twice through 40µm cell strainers and centrifuged for 10 minutes at 300g. Cells pellet were resuspended in washing solution (0.4 M mannitol, 20 mM MES, 20 mM KCl and 0.1% BSA) to a concentration of ∼1000 cells/µl and loaded on a 10X Chromium Single Cell 3’ Reagent Kit User Guide according to manufacturer protocols. Two replicates were performed with wild type and *quad* sample on each. One replicate was set to 5,000 captured cells and the second to 10,000. Libraries were sequenced using a Novaseq SP100 sequencing kit. Sequences were processed using Cell Ranger 7.0.1 and the Arabidopsis thaliana Araport11 genome annotation. Each single cell sample was processed to remove ambient RNA using SoupX(*71*) and doublet removed using DoubletFinder(*72*) using the first 15 PCA dimensions. Cell profiles having more than 5% mitochondria or chloroplast gene expression exhibiting expression of CHLOROPHYLL A/B BINDING PROTEINs were removed. Reference single-cell data matrix was obtained from GEO Omnibus series GSE123818(*73*). Cells having less than 500 UMIs, more than 25,000 UMIs, less than 500 genes, or more than 10,000 genes were removed. Genes expressed in less than 10 cells or plastid genes were removed. After filtering, we obtained 6,807, 3,722, and 3,315 cells for WT, *quad,* and reference uncut meristem, respectively (mean 6,782 UMI and 2,224 genes per cell).

The five samples were combined using Harmony(*74*) and the first 20 dimensions of the PCA. UMAP and clustering were derived using Seurat 4.1.3. Clusters with less than 10 cells were discarded. For marker analysis, we obtained previously published tissue-specific markers(*57*) and computed the average expression of these markers for each cell to get a tissue score. For the Index of Cell Identity analysis, we used the ICI algorithm as previously described(*56*), with an information threshold of 25 and a p-value cut-off of 0.05. If cells had more than one significant identity, they were assigned to the one with the lowest p-value. If cells had more than 4 significant identities, they were marked as ‘mixed.’ R scripts are provided as supplementary text. Raw single-cell data and post-processing Seurat object are deposited in GEO Omnibus GSE248204.

---

**Fig. S1.**
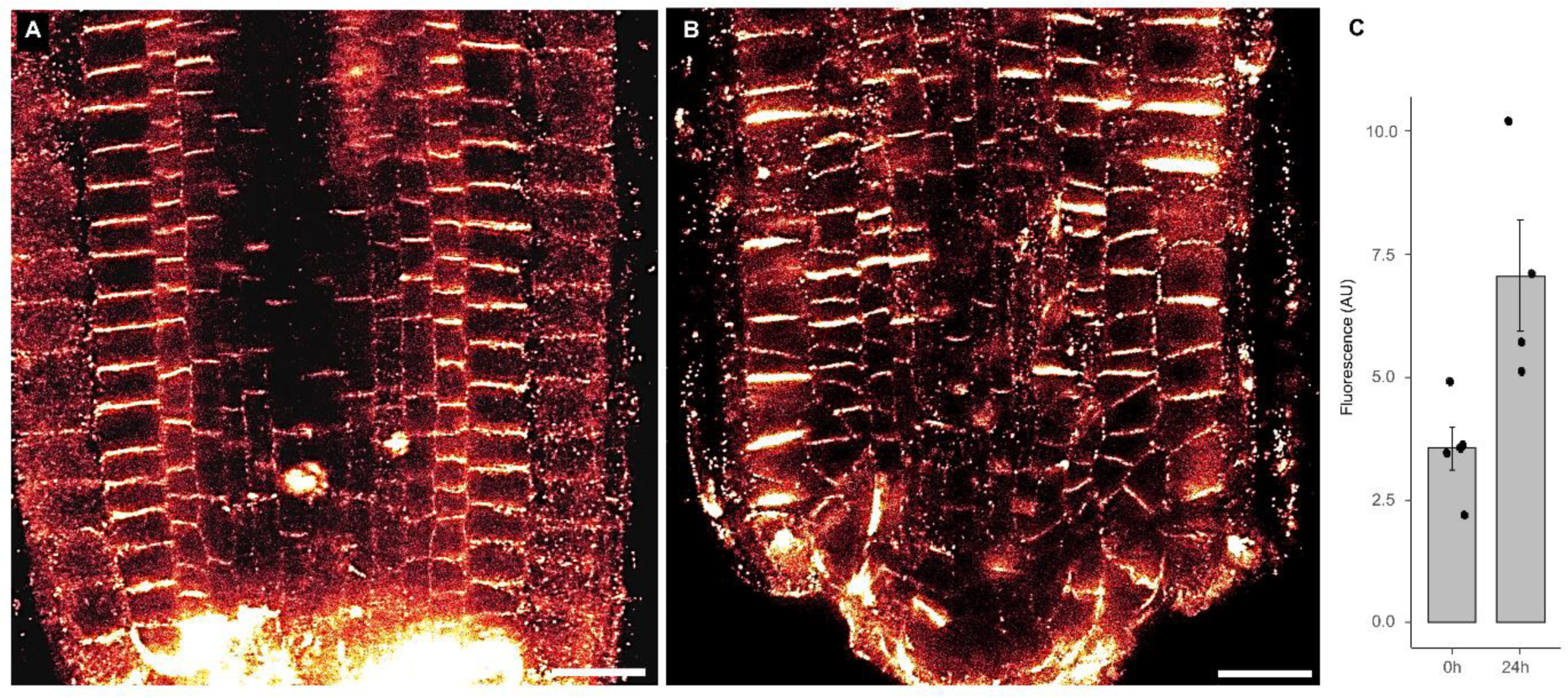
Callose localization in regeneration. **A-B)** Immunostaining using anti-callose antibodies at 0h (A) and 24h (B) after the cut (full images of zoomed-in panel in Fig. 1P-Q). **C)** Quantification of callose signal. P-val = 0.02; student’s t-test; n=5 and 4 for 0h and 24h samples, respectively. Scale bars are 25μm.

**Fig. S2.**
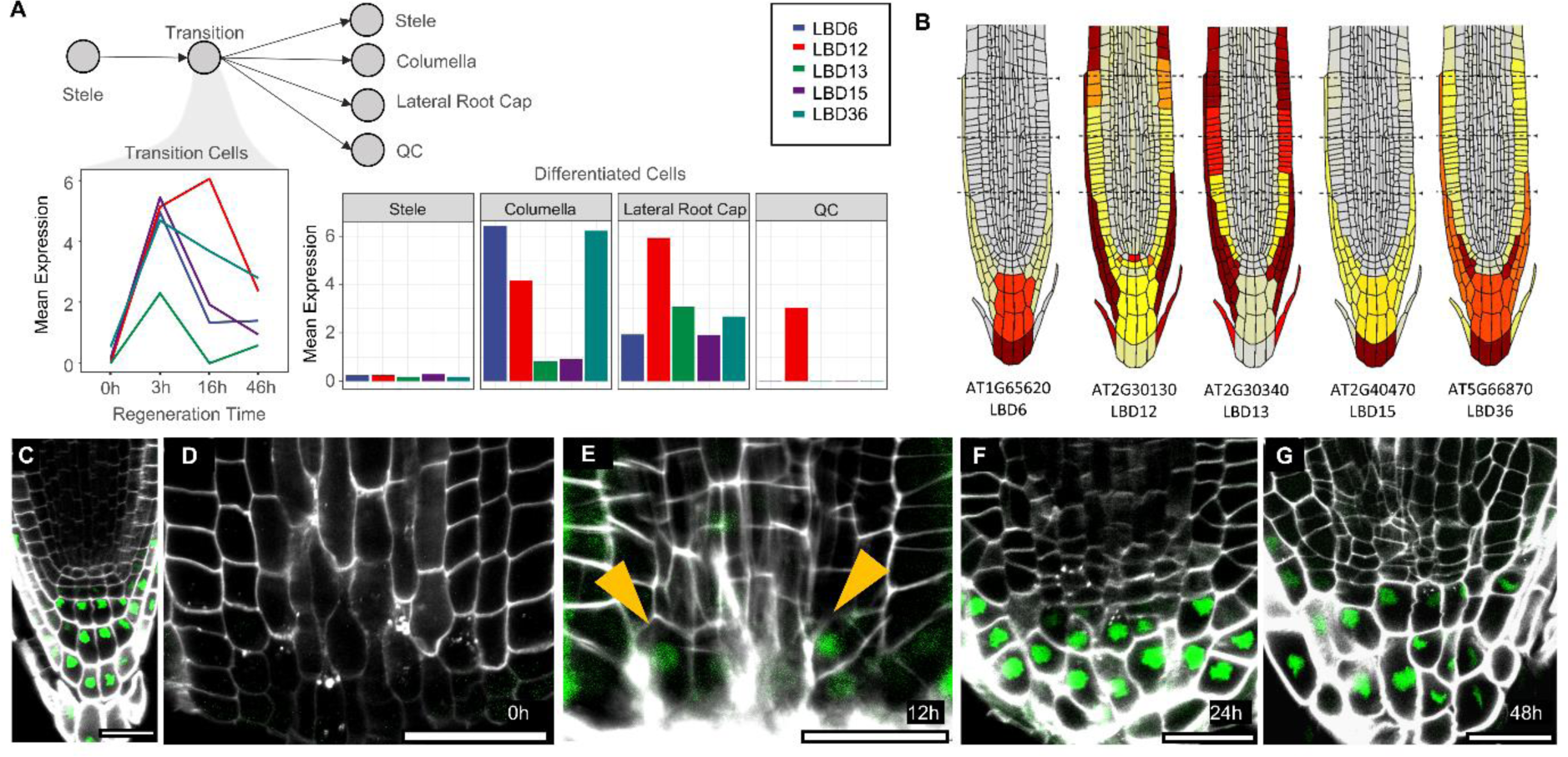
Expression patterns and overexpression of GATEKEEPER genes. **A)** Average expression of GATEKEEPER LBD in distal cells during identity transition and in mature tissues, based on data obtained from single-cell RNAseq-based expression (Efroni et al., 2016). The illustration shows the identity trajectory of cells in the stump during regeneration. **B)** Single-cell-based (www.rootatlas.com) expression patterns in intact roots. **C-G)** Confocal images of *pLBD36:NeonGreen-N7:LBD36^ter^* expression in intact root meristem (C), at 0h (D), 12h (E), 24h (F) and 48h (G) after excision. Arrowhead points to newly induced expression. Scale bars are 25μm.

**Fig. S3.**
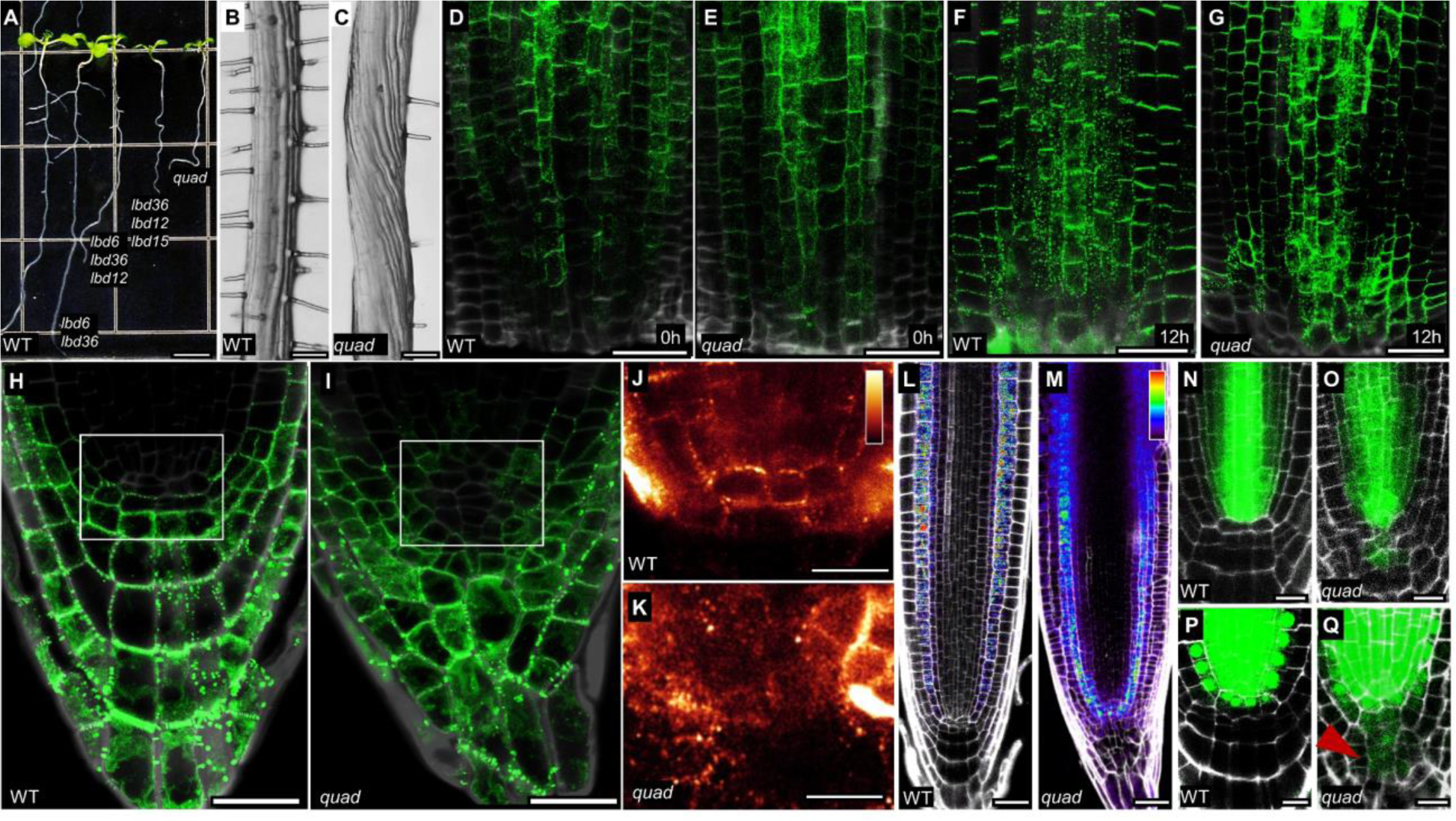
Characterization of *gatekeeper* mutant roots. **A)** 9-day-old seedling of WT and *gatekeeper* mutants. **B-C)** Stereoscope images of WT (B) and *quad* (C) roots at the differentiation zone. **D-I)** Confocal images of *35S:PDLP5-GFP* uncut root meristems (H-I), at 0h (D-E) and 12h (F-G) after excision in WT (D, F, H) or *quad* (E, G, I) backgrounds. **J-K)** Callose immunostaining in WT (J) and *quad* (K) roots stem cell niche region. **L-M)** Root meristems of WT (L) and *quad* (M) after HPTS injection. **N-Q)** Confocal images of *pSHR:ER-GFP* (N-O) and *pSHR:SHR-GFP* (P-Q) in WT (N, P) or *quad* (O, Q) backgrounds. Red arrowhead points to the ectopic presence of SHR-GFP protein. Scale bars are 500μm (A), 100μm (B-C), 25μm (D-M), and 10μm (P-Q).

**Fig. S4.**
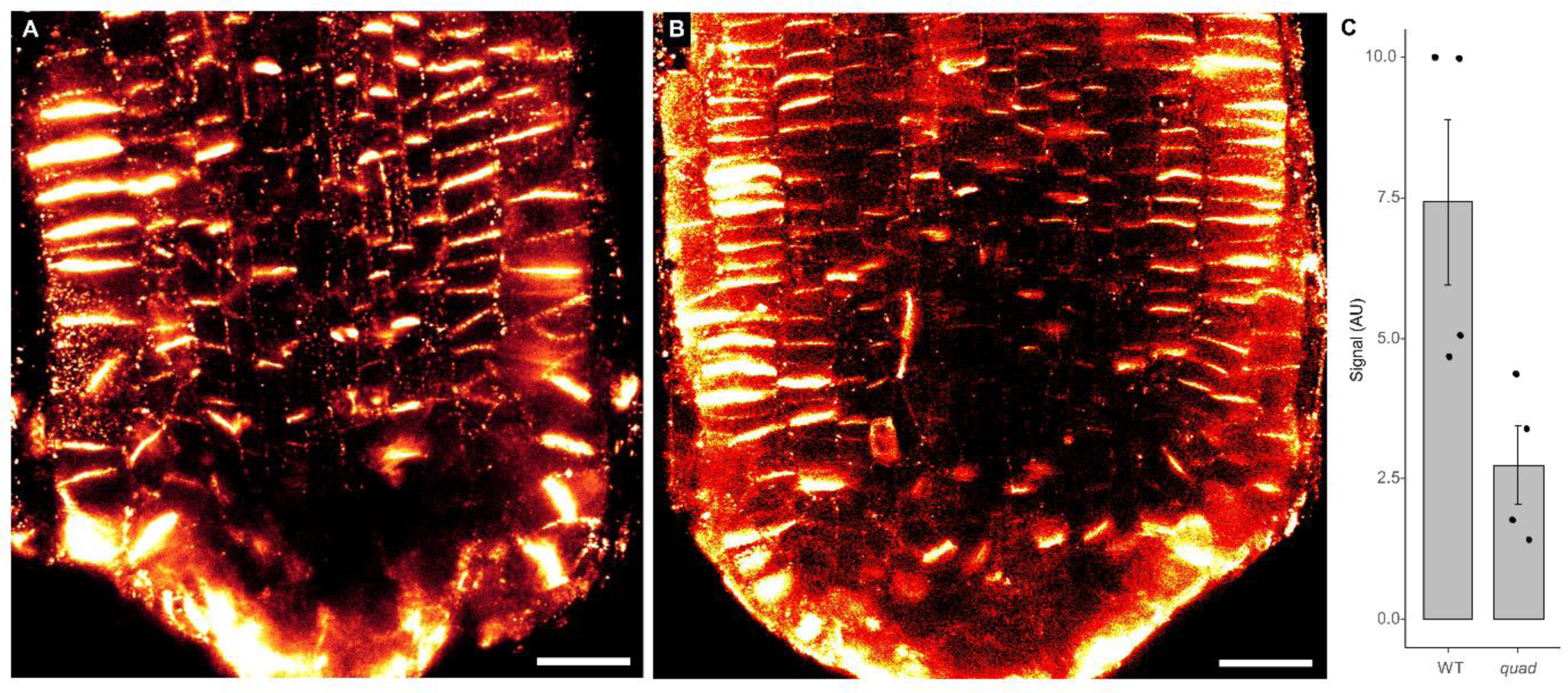
Callose localization in quad mutants. **A-B)** Immunostaining using anti-callose antibodies of WT (A) and *quad* mutants (B) at 24h after the cut (full images of zoomed-in panel in Fig. 2Q-R). **C)** Quantification of callose signal. P-val = 0.02; student’s t-test; n=4 for each sample. Scale bars are 25μm.

**Fig. S5.**
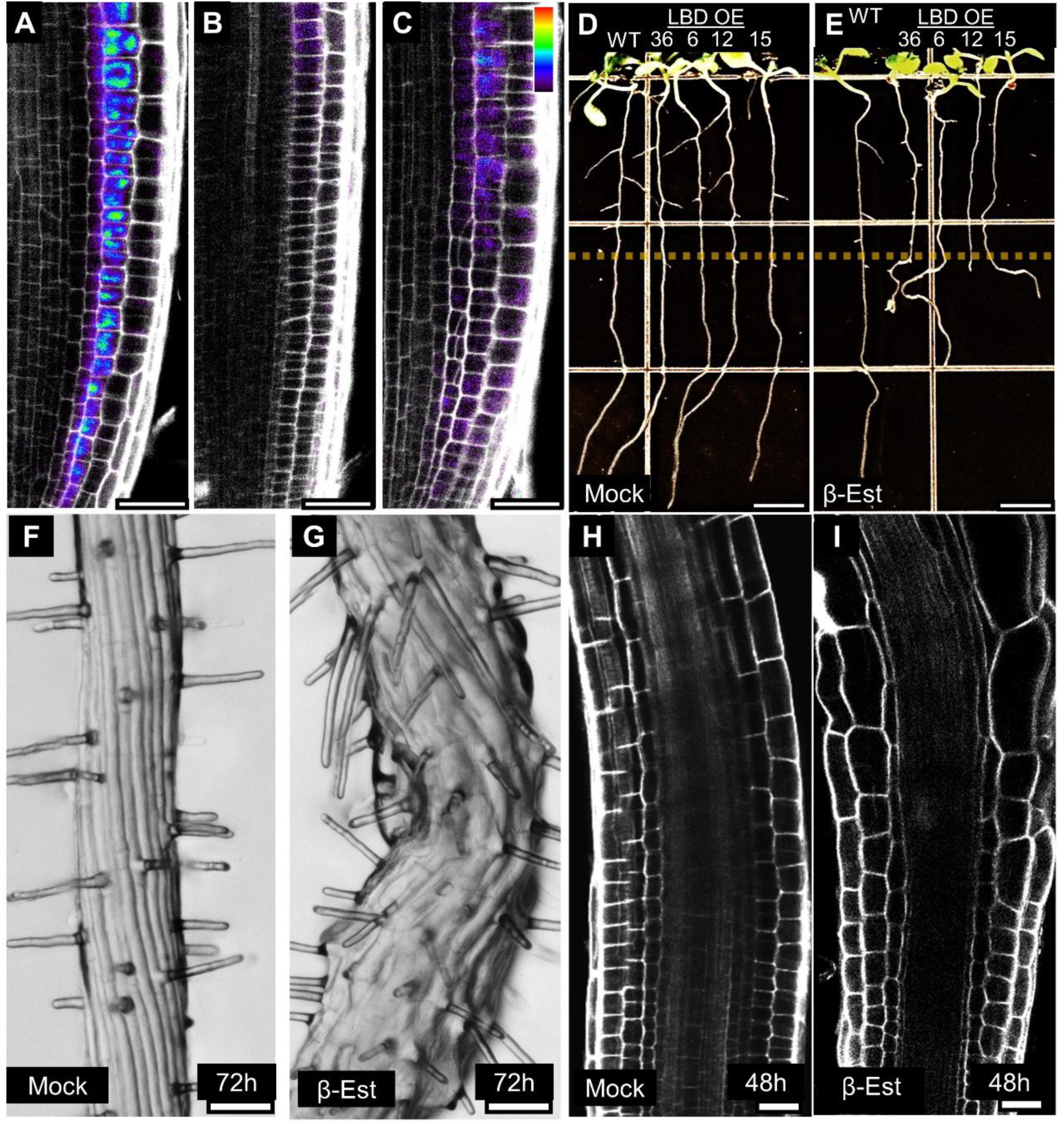
Overexpression of GATEKEEPER LBDs restricts symplastic transport and alters root development. **A-C)** Confocal images of *35:XVE lexA:LBD12* (D-E) and *35S:XVE lexA:LBD15* (F) roots treated with mock (A) or β-estradiol (B-C) for 48h, and injected with HPTS. **D-E)** 9-day-old seedling of wild-type and inducible LBD overexpression (OE) lines treated with mock (D) or β-estradiol (E) for 4 days. Dashed line marks the root length at the time of induction. **F-G)** Stereoscope images of *35:XVE-LBD36* roots treated with mock (F) or β-estradiol (G) for 72h**. H-I**) Confocal images of the transition and elongation zones of *35:XVE-LBD36* roots treated with mock (H) or β-estradiol (I) for 48h. Scale bars are 25µm (A-C, F-I) and 500µm (D-E).

**Fig. S6.**
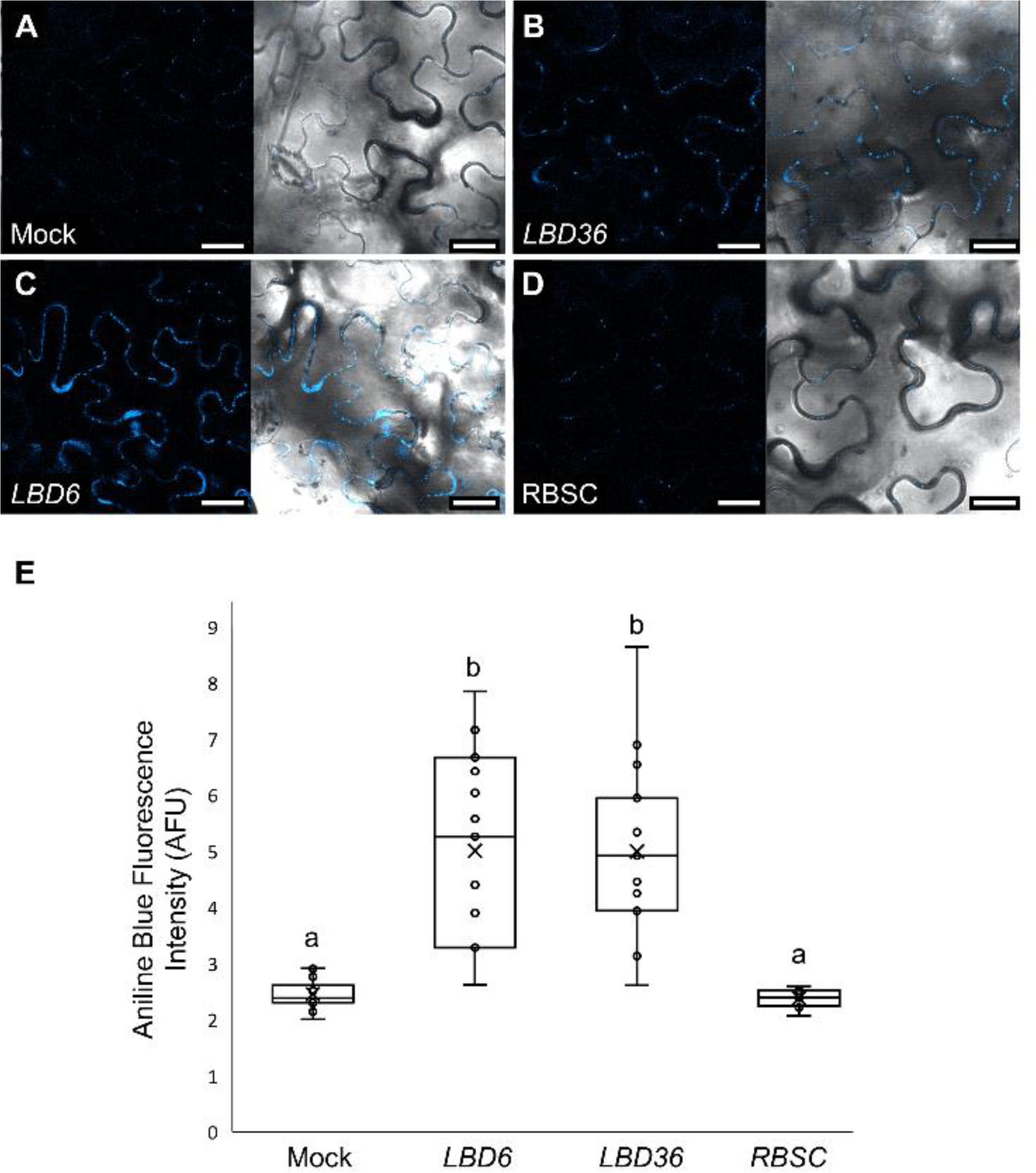
Ectopic expression of *GATEKEEPER* genes induces callose accumulation in tobacco epidermal cells. **A-D)** Aniline blue callose staining of N. benthamiana epidermal cells agroinfiltrated with empty vector (A), *35S:LBD6* (B), *35S:LBD36* (C) and *35S:RBSC* (D). **E)** Quantitative analysis of callose accumulation (Tukey’s HSD test, P-val<0.05, n ≥ 15). Scale bars are 25 μm.

**Fig. S7.**
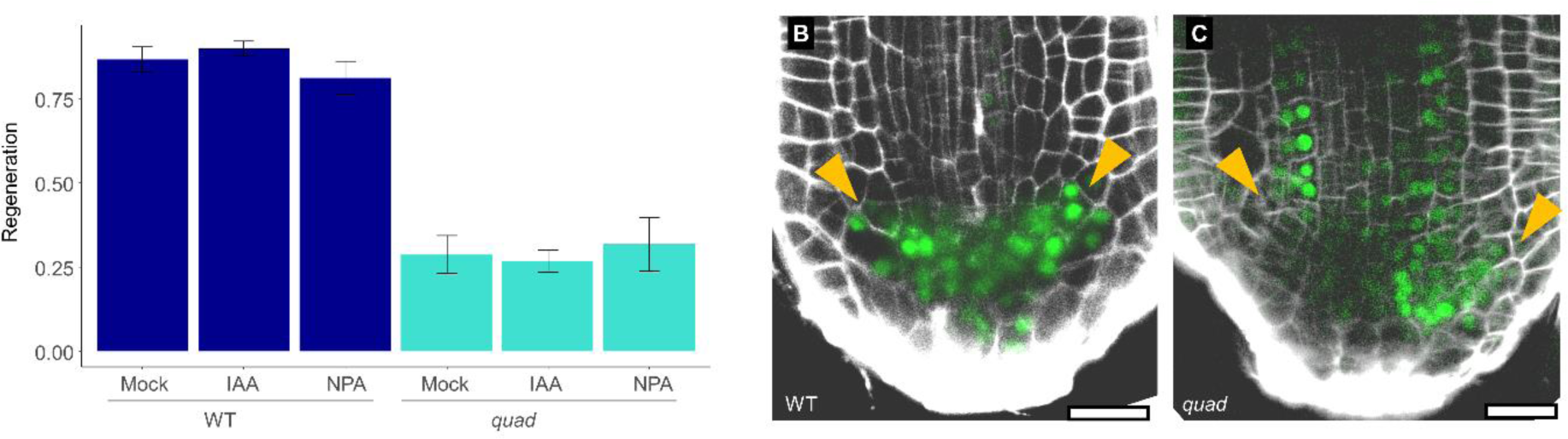
Effects of auxin and NPA treatment on regeneration in *quad* mutant. **A)** Regeneration rates in WT and *quad* treated with mock, IAA (10nM), or NPA (10µM). n=3. **B-C)** Confocal images of WT (B) and *quad* (C) root meristems at 24h after the cut, treated with 10µM NPA. NPA causes a lateral expansion of DR5 expression (marked by yellow arrowheads). Similar expansion is observed in *quad* mutants, which maintain the broad proximally expanded domain. Scale bars are 25μm.

**Fig. S8.**
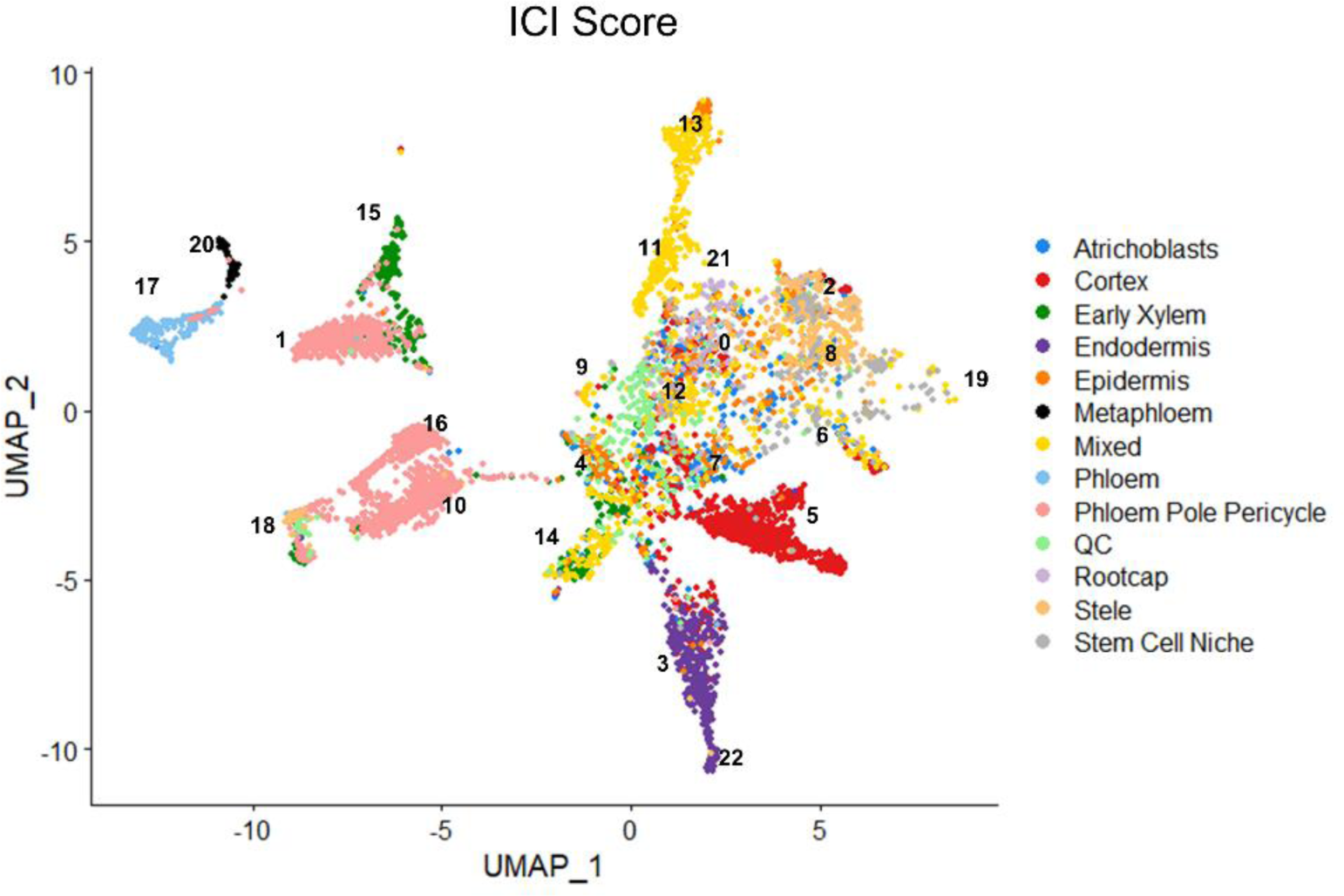
Index of Cell Identity scores for combined single-cell dataset.

**Fig. S9.**
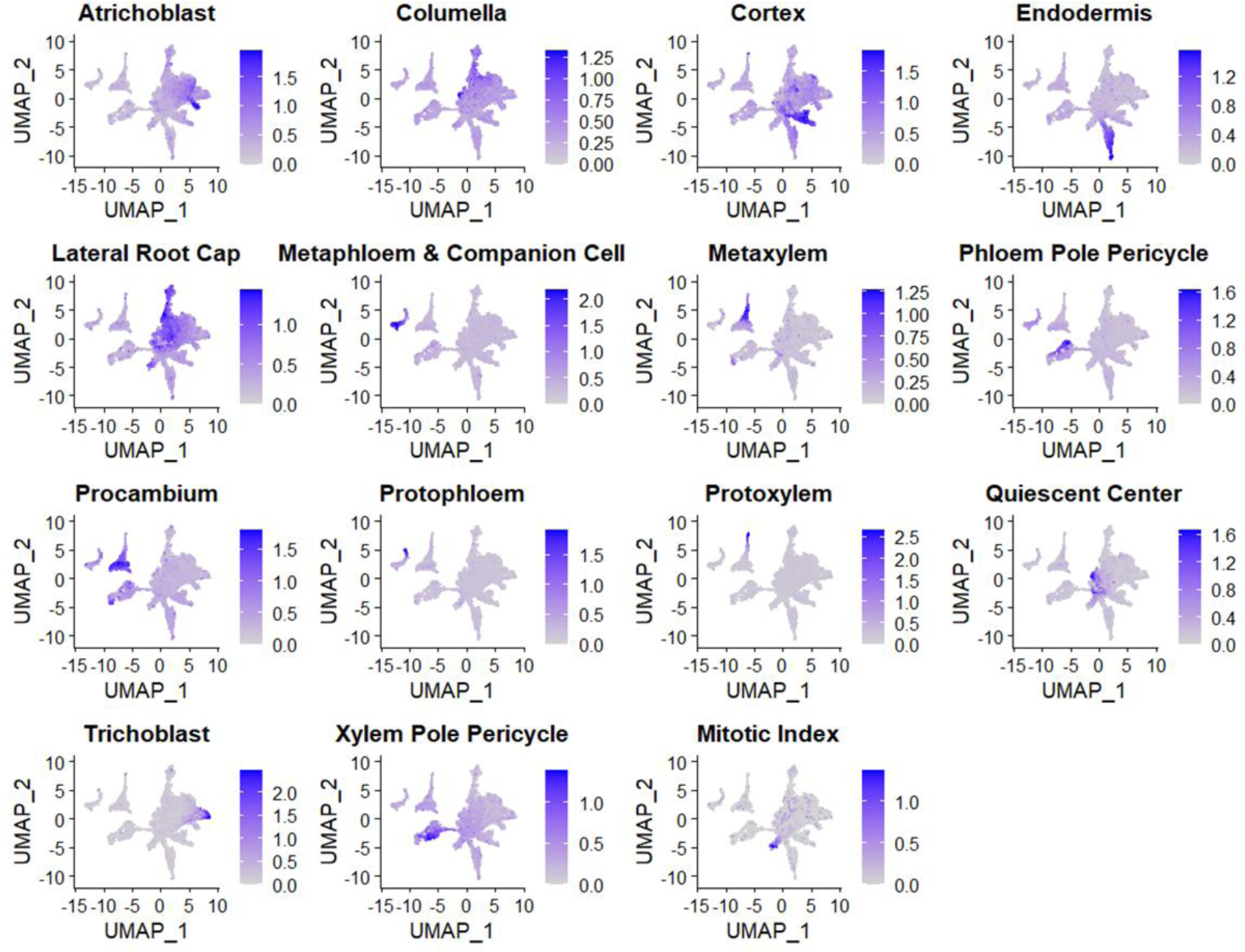
Average expression of known tissue-specific markers in single-cell dataset.

**Fig. S10.**
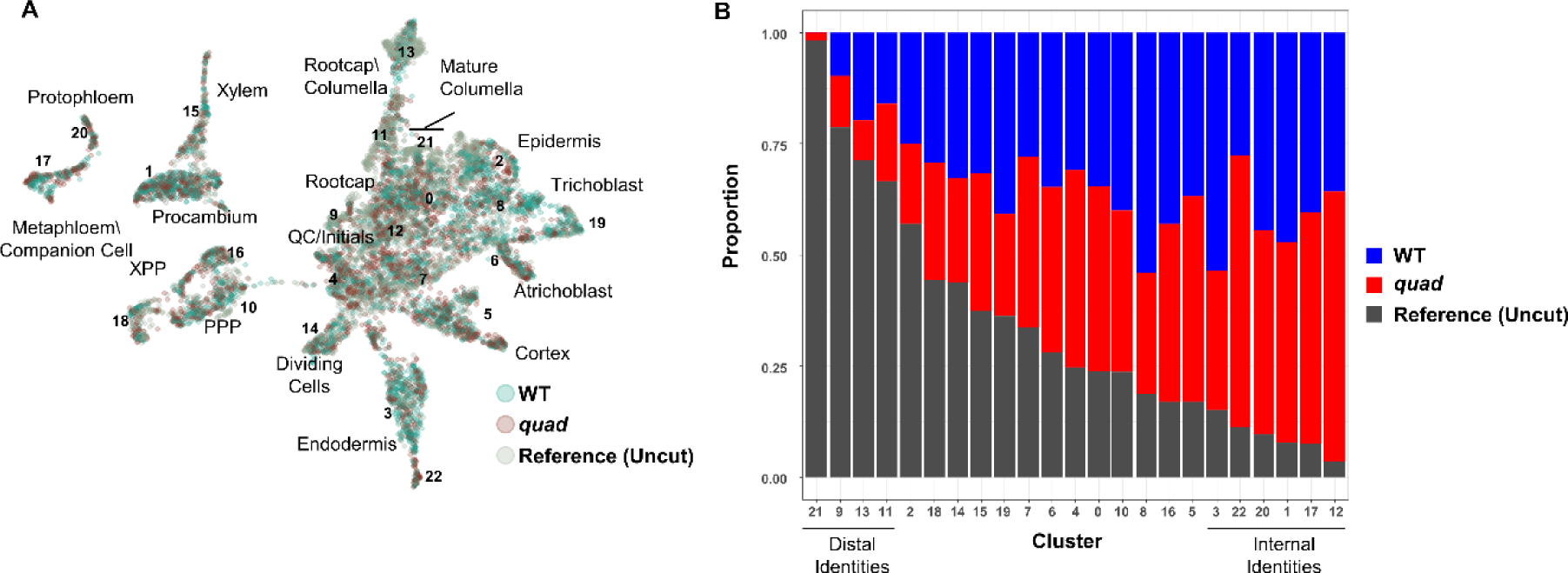
Distribution of different genotypes in single-cell clusters. **A)** UMAP of single-cell profiles used in this study, color-coded for their source. **B)** The relative representation of cells from each source in clusters.

**Table S1.** List of *gatekeeper lbd* alleles used in this study.

| Locus | Allele | Genotype | Notes |
| --- | --- | --- | --- |
| LBD6\AS2 | *lbd6-201* | g.2381_insA | Used in *lbd6 lbd36* double |
|  | *lbd6-202* | g.2381_delA | Used in *quad* |
|  | *lbd6-203* | g.2226_2233_del |  |
|  | *lbd6-204* | g.2381_2386_del |  |
| LBD12 | *lbd12-2* | g.123_739_del | Used in *lbd12 lbd15* and in *quad* |
|  | *lbd12-3* | g.773_777_del |  |
|  | *lbd12-4* | g.125_777_inv |  |
| LBD15 | *lbd15-2* | g.877_907_del | Used in *lbd12 lbd15* and in *quad* |
|  | *lbd15-3* | g.907_insT |  |
|  | *lbd15-4* | g.906_delG |  |
| LBD36 | *lbd36-2* | g.1529_insC | Used in *lbd6 lbd36* double |
|  | *lbd36-3* | g.1524_1543_del | Used in *quad* |
|  | *lbd36-4* | g.1529_1533_del |  |
|  | *lbd36-5* | g.1429_ins(23bp) |  |

**Table S2.**
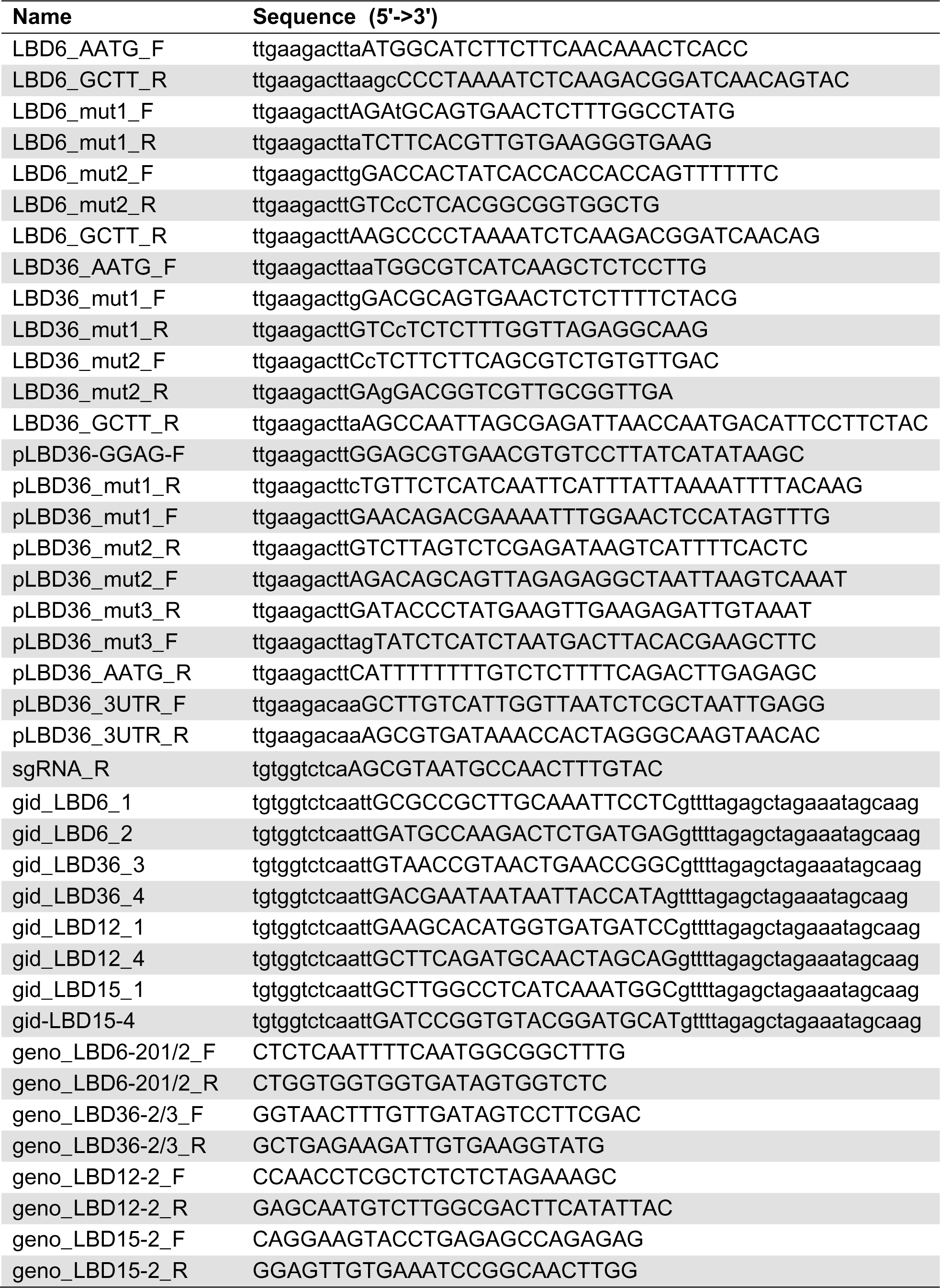
List primers used in this study.

## References

1. G. Sena, X. Wang, H.-Y. Liu, H. Hofhuis, K. D. Birnbaum, Organ regeneration does not require a functional stem cell niche in plants. Nature 457, 1150–3 (2009).

2. J. Heyman, T. Cools, B. Canher, S. Shavialenka, J. Traas, I. Vercauteren, H. Van den Daele, G. Persiau, G. De Jaeger, K. Sugimoto, L. De Veylder, The heterodimeric transcription factor complex ERF115–PAT1 grants regeneration competence. Nature Plants 2, 16165–16165 (2016).

3. R. Matosevich, I. Cohen, N. Gil-Yarom, A. Modrego, L. Friedlander-Shani, C. Verna, E. Scarpella, I. Efroni, Local auxin biosynthesis is required for root regeneration after wounding. Nature Plants 6, 1020–1030 (2020).

4. K. Durgaprasad, M. V. Roy, A. Venugopal M., A. Kareem, K. Raj, V. Willemsen, A. P. Mähönen, B. Scheres, K. Prasad, Gradient Expression of Transcription Factor Imposes a Boundary on Organ Regeneration Potential in Plants. Cell Reports 29, 453–463.e3 (2019).

5. I. Efroni, A. Mello, T. Nawy, P. Ip, R. Rahni, N. DelRose, A. Powers, R. Satija, K. D. Birnbaum, Root Regeneration Triggers an Embryo-like Sequence Guided by Hormonal Interactions. Cell 165, 1721–33 (2016).

6. Z. P. Li, A. Paterlini, M. Glavier, E. M. Bayer, Intercellular trafficking via plasmodesmata: molecular layers of complexity. Cellular and Molecular Life Sciences 78, 799–816 (2020).

7. M. Kitagawa, T. M. Tran, D. Jackson, Traveling with purpose: cell-to-cell transport of plant mRNAs. Trends in Cell Biology In press (2023).

8. K. L. Gallagher, R. Sozzani, C.-M. Lee, Intercellular Protein Movement: Deciphering the Language of Development. Annual Review of Cell and Developmental Biology 30, 207–233 (2014).

9. A. Vatén, J. Dettmer, S. Wu, Y. D. Stierhof, S. Miyashima, S. R. Yadav, C. J. Roberts, A. Campilho, V. Bulone, R. Lichtenberger, S. Lehesranta, A. P. Mähönen, J. Y. Kim, E. Jokitalo, N. Sauer, B. Scheres, K. Nakajima, A. Carlsbecker, K. L. Gallagher, Y. Helariutta, Callose Biosynthesis Regulates Symplastic Trafficking during Root Development. Developmental Cell 21, 1144–1155 (2011).

10. Y. Helariutta, H. Fukaki, J. Wysocka-Diller, K. Nakajima, J. Jung, G. Sena, M.-T. Hauser, P. N. Benfey, The SHORT-ROOT Gene Controls Radial Patterning of the Arabidopsis Root through Radial Signaling. Cell 101, 555–567 (2000).

11. K. Nakajima, G. Sena, T. Nawy, P. N. Benfey, Intercellular movement of the putative transcription factor SHR in root patterning. Nature 413, 307–311 (2001).

12. Z. Spiegelman, C.-M. Lee, K. L. Gallagher, KinG Is a Plant-Specific Kinesin That Regulates Both Intra- and Intercellular Movement of SHORT-ROOT. Plant Physiology 176, 392–405 (2018).

13. K. L. Gallagher, P. N. Benfey, Both the conserved GRAS domain and nuclear localization are required for SHORT-ROOT movement. The Plant Journal 57, 785–797 (2009).

14. A. Vatén, J. Dettmer, S. Wu, Y.-D. Stierhof, S. Miyashima, S. R. Yadav, C. J. Roberts, A. Campilho, V. Bulone, R. Lichtenberger, S. Lehesranta, A. P. Mähönen, J.-Y. Kim, E. Jokitalo, N. Sauer, B. Scheres, K. Nakajima, A. Carlsbecker, K. L. Gallagher, Y. Helariutta, Callose Biosynthesis Regulates Symplastic Trafficking during Root Development. Developmental Cell 21, 1144–1155 (2011).

15. G. Sena, J. W. Jung, P. N. Benfey, A broad competence to respond to SHORT ROOT revealed by tissue-specific ectopic expression. Development 131, 2817–2826 (2004).

16. S. Wu, R. O’Lexy, M. Xu, Y. Sang, X. Chen, Q. Yu, K. L. Gallagher, Symplastic signaling instructs cell division, cell expansion, and cell polarity in the ground tissue of *Arabidopsis thaliana* roots. Proc. Natl. Acad. Sci. U.S.A. 113, 11621–11626 (2016).

17. K. Koizumi, T. Hayashi, S. Wu, K. L. Gallagher, The SHORT-ROOT protein acts as a mobile, dose-dependent signal in patterning the ground tissue. Proc. Natl. Acad. Sci. U.S.A. 109, 13010– 13015 (2012).

18. S. Sabatini, D. Beis, H. Wolkenfelt, J. Murfett, T. Guilfoyle, J. Malamy, P. Benfey, O. Leyser, N. Bechtold, P. Weisbeek, B. Scheres, An Auxin-Dependent Distal Organizer of Pattern and Polarity in the Arabidopsis Root. Cell 99, 463–472 (1999).

19. L. R. Band, Auxin fluxes through plasmodesmata. New Phytologist 231, 1686–1692 (2021).

20. M. Adamowski, J. Friml, PIN-Dependent Auxin Transport: Action, Regulation, and Evolution. The Plant Cell 27, 20–32 (2015).

21. P. Mehra, B. K. Pandey, D. Melebari, J. Banda, N. Leftley, V. Couvreur, J. Rowe, M. Anfang, H. De Gernier, E. Morris, C. J. Sturrock, S. J. Mooney, R. Swarup, C. Faulkner, T. Beeckman, R. P. Bhalerao, E. Shani, A. M. Jones, I. C. Dodd, R. E. Sharp, A. Sadanandom, X. Draye, M. J. Bennett, Hydraulic flux–responsive hormone redistribution determines root branching. Science 378, 762–768 (2022).

22. R. Sager, X. Wang, K. Hill, B. C. Yoo, J. Caplan, A. Nedo, T. Tran, M. J. Bennett, J. Y. Lee, Auxin-dependent control of a plasmodesmal regulator creates a negative feedback loop modulating lateral root emergence. Nature Communications 11, 1–10 (2020).

23. N. M. Linh, E. Scarpella, Leaf vein patterning is regulated by the aperture of plasmodesmata intercellular channels. PLoS Biol 20, e3001781 (2022).

24. Y. Liu, M. Xu, N. Liang, Y. Zheng, Q. Yu, S. Wu, Symplastic communication spatially directs local auxin biosynthesis to maintain root stem cell niche in *Arabidopsis*. Proc. Natl. Acad. Sci. U.S.A. 114, 4005–4010 (2017).

25. P. L. H. Rinne, C. van der Schoot, Symplasmic fields in the tunica of the shoot apical meristem coordinate morphogenetic events. Development 125, 1477–1485 (1998).

26. E. Bayer, C. Thomas, A. Maule, Symplastic domains in the Arabidopsis shoot apical meristem correlate with PDLP1 expression patterns. Plant Signal Behav 3, 853–855 (2008).

27. Y. Benitez-Alfonso, C. Faulkner, A. Pendle, S. Miyashima, Y. Helariutta, A. Maule, Symplastic Intercellular Connectivity Regulates Lateral Root Patterning. Developmental Cell 26, 136–147 (2013).

28. R. Gaudioso-Pedraza, M. Beck, L. Frances, P. Kirk, C. Ripodas, A. Niebel, G. E. D. Oldroyd, Y. Benitez-Alfonso, F. de Carvalho-Niebel, Callose-Regulated Symplastic Communication Coordinates Symbiotic Root Nodule Development. Current Biology 28, 3562–3577.e6 (2018).

29. I. Kim, K. Kobayashi, E. Cho, P. C. Zambryski, Subdomains for transport via plasmodesmata corresponding to the apical–basal axis are established during *Arabidopsis* embryogenesis. Proc. Natl. Acad. Sci. U.S.A. 102, 11945–11950 (2005).

30. K. J. Oparka, Plasmodesmata (Blackwell publ, Oxford, 2005) *Annual plant reviews*.

31. R. Stadler, C. Lauterbach, N. Sauer, Cell-to-cell movement of green fluorescent protein reveals post-phloem transport in the outer integument and identifies symplastic domains in Arabidopsis seeds and embryos. Plant Physiology 139, 701–712 (2005).

32. K. Ehlers, H. Binding, R. Kollmann, The formation of symplasmic domains by plugging of plasmodesmata: a general event in plant morphogenesis? Protoplasma 209, 181–192 (1999).

33. K. Godel-Jedrychowska, K. Kulinska-Lukaszek, A. Horstman, M. Soriano, M. Li, K. Malota, K. Boutilier, E. U. Kurczynska, Symplasmic isolation marks cell fate changes during somatic embryogenesis. Journal of Experimental Botany 71, 2612–2628 (2020).

34. T. Zhu, W. J. Lucas, T. L. Rost, Directional cell-to-cell communication in the Arabidopsis root apical meristem I. An ultrastructural and functional analysis. Protoplasma 203, 35–47 (1998).

35. J. D. Petit, Z. P. Li, W. J. Nicolas, M. S. Grison, E. M. Bayer, Dare to change, the dynamics behind plasmodesmata-mediated cell-to-cell communication. Current Opinion in Plant Biology 53, 80– 89 (2020).

36. J. O. Brunkard, P. C. Zambryski, Plasmodesmata enable multicellularity: new insights into their evolution, biogenesis, and functions in development and immunity. Current Opinion in Plant Biology 35, 76–83 (2017).

37. S. Tylewicz, A. Petterle, S. Marttila, P. Miskolczi, A. Azeez, R. K. Singh, J. Immanen, N. Mähler, T. R. Hvidsten, D. M. Eklund, J. L. Bowman, Y. Helariutta, R. P. Bhalerao, Photoperiodic control of seasonal growth is mediated by ABA acting on cell-cell communication. Science 360, 212–215 (2018).

38. A. Imlau, E. Truernit, N. Sauer, Cell-to-cell and long-distance trafficking of the green fluorescent protein in the phloem and symplastic unloading of the protein into sink tissues. The Plant Cell 11, 309–322 (1999).

39. J. Y. Lee, X. Wang, W. Cui, R. Sager, S. Modla, K. Czymmek, B. Zybaliov, K. Van Wijk, C. Zhang, H. Lu, V. Lakshmanana, A plasmodesmata-localized protein mediates crosstalk between cell-to-cell communication and innate immunity in Arabidopsis. Plant Cell 23, 3353–3373 (2011).

40. Y. Benitez-Alfonso, C. Faulkner, A. Pendle, S. Miyashima, Y. Helariutta, A. Maule, Symplastic Intercellular Connectivity Regulates Lateral Root Patterning. Developmental Cell 26, 136–147 (2013).

41. E. M. Bell, W. Lin, A. Y. Husbands, L. Yu, V. Jaganatha, B. Jablonska, A. Mangeon, M. M. Neff, T. Girke, P. S. Springer, Arabidopsis LATERAL ORGAN BOUNDARIES negatively regulates brassinosteroid accumulation to limit growth in organ boundaries. Proceedings of the National Academy of Sciences 109, 21146–21151 (2012).

42. W. Lin, B. Shuai, P. S. Springer, The Arabidopsis LATERAL ORGAN BOUNDARIES–Domain Gene ASYMMETRIC LEAVES2 Functions in the Repression of KNOX Gene Expression and in Adaxial-Abaxial Patterning. The Plant Cell 15, 2241–2252 (2003).

43. Y. Machida, T. Suzuki, M. Sasabe, H. Iwakawa, S. Kojima, C. Machida, Arabidopsis ASYMMETRIC LEAVES2 (AS2): roles in plant morphogenesis, cell division, and pathogenesis. J Plant Res 135, 3–14 (2022).

44. Y. Okushima, H. Fukaki, M. Onoda, A. Theologis, M. Tasaka, ARF7 and ARF19 regulate lateral root formation via direct activation of LBD/ASL genes in Arabidopsis. The Plant Cell 19, 118–30 (2007).

45. T. Goh, K. Toyokura, N. Yamaguchi, Y. Okamoto, T. Uehara, S. Kaneko, Y. Takebayashi, H. Kasahara, Y. Ikeyama, Y. Okushima, K. Nakajima, T. Mimura, M. Tasaka, H. Fukaki, Lateral root initiation requires the sequential induction of transcription factors LBD16 and PUCHI in Arabidopsis thaliana. New Phytologist 224, 749–760 (2019).

46. M. Omary, N. Gil-Yarom, C. Yahav, E. Steiner, I. Efroni, A conserved superlocus regulates above- and belowground root initiation. Science 375, eabf4368 (2022).

47. T. Goh, S. Joi, T. Mimura, H. Fukaki, The establishment of asymmetry in Arabidopsis lateral root founder cells is regulated by LBD16/ASL18 and related LBD/ASL proteins. Development 139, 883–893 (2012).

48. L. Ye, X. Wang, M. Lyu, R. Siligato, G. Eswaran, L. Vainio, T. Blomster, Cytokinins initiate secondary growth in the Arabidopsis root through a set of LBD genes. Current Biology 31, 3365–3373 (2021).

49. E. Bortiri, G. Chuck, E. Vollbrecht, T. Rocheford, R. Martienssen, S. Hake, ramosa2 Encodes a LATERAL ORGAN BOUNDARY Domain Protein That Determines the Fate of Stem Cells in Branch Meristems of Maize. The Plant Cell 18, 574–585 (2006).

50. L. Borghi, M. Bureau, R. Simon, Arabidopsis JAGGED LATERAL ORGANS is expressed in boundaries and coordinates KNOX and PIN activity. The Plant Cell 19, 1795–808 (2007).

51. M. Fan, C. Xu, K. Xu, Y. Hu, LATERAL ORGAN BOUNDARIES DOMAIN transcription factors direct callus formation in Arabidopsis regeneration. Cell Research 22, 1169–1180 (2012).

52. M. I. Rast, R. Simon, Arabidopsis JAGGED LATERAL ORGANS Acts with ASYMMETRIC LEAVES2 to Coordinate KNOX and PIN Expression in Shoot and Root Meristems. The Plant Cell 24, 2917– 2933 (2012).

53. C. Cho, E. Jeon, S. K. Pandey, S. H. Ha, J. Kim, LBD13 positively regulates lateral root formation in Arabidopsis. Planta 249, 1251–1258 (2019).

54. A. S. Chanderbali, F. He, P. S. Soltis, D. E. Soltis, Out of the water: Origin and diversification of the LBD gene family. Molecular Biology and Evolution 32, 1996–2000 (2015).

55. J. R. Wendrich, B. Yang, N. Vandamme, K. Verstaen, W. Smet, C. Van de Velde, M. Minne, B. Wybouw, E. Mor, H. E. Arents, J. Nolf, J. Van Duyse, G. Van Isterdael, S. Maere, Y. Saeys, B. De Rybel, Vascular transcription factors guide plant epidermal responses to limiting phosphate conditions. Science 370, eaay4970 (2020).

56. I. Efroni, P.-L. Ip, T. Nawy, A. Mello, K. D. Birnbaum, Quantification of cell identity from single-cell gene expression profiles. Genome biology 16, 1–12 (2015).

57. R. Shahan, C.-W. Hsu, T. M. Nolan, B. J. Cole, I. W. Taylor, L. Greenstreet, S. Zhang, A. Afanassiev, A. H. C. Vlot, G. Schiebinger, P. N. Benfey, U. Ohler, A single-cell Arabidopsis root atlas reveals developmental trajectories in wild-type and cell identity mutants. Developmental Cell 57, 543–560.e9 (2022).

58. M. Ackerman-Lavert, Y. Fridman, R. Matosevich, H. Khandal, L. Friedlander-Shani, K. Vragović, R. Ben El, G. Horev, D. Tarkowská, I. Efroni, S. Savaldi-Goldstein, Auxin requirements for a meristematic state in roots depend on a dual brassinosteroid function. Current Biology 31, 4462–4472.e6 (2021).

59. Y. Matsumura, H. Iwakawa, Y. Machida, C. Machida, Characterization of genes in the ASYMMETRIC LEAVES2/LATERAL ORGAN BOUNDARIES (AS2/LOB) family in Arabidopsis thaliana, and functional and molecular comparisons between AS2 and other family members. The Plant Journal 58, 525–537 (2009).

60. A. Mangeon, W. Lin, P. S. Springer, Functional divergence in the Arabidopsis LOB-domain gene family. Plant Signal Behav 7, 1544–1547 (2012).

61. K. Schiessl, J. L. S. Lilley, T. Lee, I. Tamvakis, W. Kohlen, P. C. Bailey, A. Thomas, J. Luptak, K. Ramakrishnan, M. D. Carpenter, K. S. Mysore, J. Wen, S. Ahnert, V. A. Grieneisen, G. E. D. Oldroyd, NODULE INCEPTION Recruits the Lateral Root Developmental Program for Symbiotic Nodule Organogenesis in Medicago truncatula. Current Biology 29, 3657–3668.e5 (2019).

62. T. Soyano, Y. Shimoda, M. Kawaguchi, M. Hayashi, A shared gene drives lateral root development and root nodule symbiosis pathways in Lotus. Science 366, 1021–1023 (2019).

63. Y. Helariutta, H. Fukaki, J. Wysocka-Diller, K. Nakajima, J. Jung, G. Sena, M. T. Hauser, P. N. Benfey, The SHORT-ROOT gene controls radial patterning of the Arabidopsis root through radial signaling. Cell 101, 555–67 (2000).

64. M. G. Heisler, C. Ohno, P. Das, P. Sieber, G. V. Reddy, J. A. Long, E. M. Meyerowitz, Patterns of Auxin Transport and Gene Expression during Primordium Development Revealed by Live Imaging of the Arabidopsis Inflorescence Meristem. Current Biology 15, 1899–1911 (2005).

65. B. C. W. Crawford, J. Sewell, G. Golembeski, C. Roshan, J. A. Long, M. F. Yanofsky, Genetic control of distal stem cell fate within root and embryonic meristems. Science 347, 655–659 (2015).

66. S. Werner, C. Engler, E. Weber, R. Gruetzner, S. Marillonnet, Fast track assembly of multigene constructs using golden gate cloning and the MoClo system. Bioengineered Bugs 3, 38–43 (2012).

67. J. Zuo, Q. W. Niu, N. H. Chua, An estrogen receptor-based transactivator XVE mediates highly inducible gene expression in transgenic plants. Plant Journal 24, 265–273 (2000).

68. D. Kurihara, Y. Mizuta, Y. Sato, T. Higashiyama, ClearSee: a rapid optical clearing reagent for whole-plant fluorescence imaging. Development 142, 4168–79 (2015).

69. M. Heinlein, Ed., Plasmodesmata: Methods and Protocols (Springer New York, New York, NY, 2015; https://link.springer.com/10.1007/978-1-4939-1523-1)vol. 1217 of *Methods in Molecular Biology*.

70. T. Pasternak, O. Tietz, K. Rapp, M. Begheldo, R. Nitschke, B. Ruperti, K. Palme, Protocol: an improved and universal procedure for whole-mount immunolocalization in plants. Plant Methods 11, 50 (2015).

71. M. D. Young, S. Behjati, SoupX removes ambient RNA contamination from droplet-based single-cell RNA sequencing data. GigaScience 9, giaa151 (2020).

72. C. S. McGinnis, L. M. Murrow, Z. J. Gartner, DoubletFinder: Doublet Detection in Single-Cell RNA Sequencing Data Using Artificial Nearest Neighbors. Cell Systems 8, 329–337.e4 (2019).

73. T. Denyer, X. Ma, S. Klesen, E. Scacchi, K. Nieselt, M. C. P. Timmermans, Spatiotemporal Developmental Trajectories in the Arabidopsis Root Revealed Using High-Throughput Single-Cell RNA Sequencing. Developmental Cell 48, 840–852.e5 (2019).

74. I. Korsunsky, N. Millard, J. Fan, K. Slowikowski, F. Zhang, K. Wei, Y. Baglaenko, M. Brenner, P. ru Loh, S. Raychaudhuri, Fast, sensitive and accurate integration of single-cell data with Harmony. Nature Methods 16, 1289–1296 (2019).

